# Rapid specification of human pluripotent stem cells to functional astrocytes

**DOI:** 10.1101/2022.08.25.505166

**Authors:** B. Lendemeijer, M. Unkel, B. Mossink, S. Hijazi, S.G. Sampedro, G. Shpak, D.E. Slump, M.C.G.N. van den Hout, W.F.J. van IJcken, E.M.J. Bindels, W.J.G. Hoogendijk, N. Nadif Kasri, F.M.S. de Vrij, S.A. Kushner

## Abstract

Astrocytes are essential for the formation and maintenance of neural networks through metabolic support, facilitation of synaptic function, and optimization of electrophysiological activity. However, a major technical challenge for investigating astrocyte function and disease-related pathophysiology has been the limited ability to obtain functional human astrocytes. Here we present a novel method to efficiently differentiate human pluripotent stem cell (hPSC)-derived neural progenitors to functional astrocytes in 28 days using a culture medium containing leukemia inhibitory factor (LIF) and bone morphogenetic protein 4 (BMP4). This approach yields highly pure populations of astrocytes expressing canonical astrocyte markers, which we confirmed by immunofluorescence, flow cytometry and RNA sequencing. Human PSC-derived astrocytes efficiently buffer glutamate and robustly support neural network activity. Co-cultures of hPSC-derived astrocytes and neurons on multi-electrode arrays generated robust network activity within 2 days and synchronous network bursts after 6 days. Whole cell patch-clamp recordings revealed an increased frequency of postsynaptic currents in human hPSC-derived neurons co-cultured with hPSC-derived versus primary rodent astrocytes, consistent with a corresponding increase in synapse density. Furthermore, hPSC-derived astrocytes retained their hominid morphology when transplanted into a mouse brain. In conclusion, we present a novel protocol to obtain functional astrocytes from human pluripotent stem cells, providing a platform for investigating human astrocyte function and neuronal-glial interactions.

## INTRODUCTION

Astrocytes are crucial for proper brain functioning and no longer considered to merely provide structural support for neurons^1^. Astrocytes provide neurons with a critical source of metabolites^2^, regulate blood flow^3^, maintain the blood-brain barrier^4^, facilitate synapse formation^5^ and influence neuronal network activity^6^. Astrocyte morphological complexity is one of the distinguishing features between the human and rodent brain^7^. Moreover, increasing evidence has highlighted robust functional differences between rodent and human astrocytes – human astrocytes propagate calcium waves at higher speed^8^ and more efficiently promote synaptogenesis^9^.

Human pluripotent stem cell (hPSC)-derived neural cells provide the opportunity to study the development of the human brain in vitro^10^. Currently, there are multiple protocols to differentiate human stem cells into neurons^11,12,13^. A widely-adopted protocol that yields a pure culture of neurons through forced *Ngn2*-overexpression^14^ requires co-culturing with astrocytes to ensure neuronal survival and maturation^15^. The currently available options for astrocyte co-culture with hPSC-derived neurons are either primary rodent astrocytes, the current golden-standard in the field, or human pluripotent stem cell-derived astrocytes. However, using rodent astrocytes can be undesirable due to the genomic and functional differences between human and rodent cells. Currently available protocols for differentiation of hPSCs into astrocytes have significant limitations, including contamination of other cell types^16^, very long differentiation periods^17^, and the inability to support functional neuronal maturation^18,19,15,20^. We have previously described protocols to differentiate iPSCs to NPCs, which can survive multiple freeze/thaw cycles, and subsequently can be developed into mature neural networks^12^. Here, we present a novel protocol to differentiate hPSC-derived NPCs into functional cortical astrocytes that promote neuronal maturation and network formation, eliminating the need for their rodent counterparts in human neural co-culture systems. Astrocytes differentiated using this protocol express the canonical astrocytic markers, integrate into the mouse brain after transplantation, buffer synaptic glutamate and promote neuronal maturation and function.

## METHODS

### Astrocyte differentiation

We validated our protocol in 4 different human pluripotent stem cell (hPSC) lines: 3 iPSC lines [WTC-11 Coriell #GM25256 (iPS1) and 2 in-house control lines^21^ (male, age 57 (iPS2), female, age 54 (iPS3)] and an embryonic stem cell (ESC) line (SA001 hESC). Pluripotent stem cells were differentiated to NPCs as previously described^12^ with slight modifications (**Supplementary Figure 1**). All cells were maintained in an incubator at 37 °C/5%CO_2_. HPSCs were expanded in hES medium (**Table 1**) on a feeder layer of mouse embryonic fibroblasts. A 60-70% confluent 6-well plate of undifferentiated hPSC colonies was lifted from the feeder layer using collagenase (Thermo Fisher, 17104019). Colonies were transferred to a 10 cm dish containing hES medium without fibroblast growth factor on a shaker (+/− 50 RPM). On day 3, the medium was changed to neural induction medium (**Table 1**) and refreshed every other day. After 7 days in suspension, EBs were collected and seeded on laminin coated dishes (20 μg/ml (Sigma, L2020)). On day 14 the medium was switched to NPC medium (**Table 1**). Cells were passaged 1:4 every week using collagenase and a cell lifter. After passage 3, NPC cultures were purified using fluorescence-activated cell sorting (FACS)^22,23^. NPCs were detached from the culture plate using ACCUTASE™ (Stemcell technologies, 07920) and resuspended into a single cell solution. CD184^+^/CD44^−^/CD271^−^/CD24^+^ cells were collected using a FACSaria III (BD bioscience) and expanded. To obtain hPSC-derived astrocytes, NPCs, passage 5-10, were passaged 1:4 using ACCUTASE™ (Stemcell technologies, 07920) when confluent and subsequently grown in Astrocyte medium containing leukemia inhibitory factor (LIF) and bone morphogenetic protein 4 (BMP4) for 4 weeks (**Table 1, Figure 1A**). Cells were grown on laminin coated dishes (20 μg/ml (Sigma, L2020)) and passaged 1:4 when confluent. The passaging ratio was adapted to the proliferation rate that gradually slows down at later stages of differentiation. During the first 2 weeks of the protocol a substantial amount of cell death was observed. Following the 4-week differentiation period, astrocytes could be maintained for at least an additional 6 weeks.

**Table 1:** Overview of media and reagents used.

| Name | Reagents | Manufacturer, catalogue number |
| --- | --- | --- |
| hES Medium | Advanced DMEM/F12<br>20% HyClone Fetal Bovine Serum<br>1% MEM-NEAA<br>7 nl/ml β-mercaptoethanol<br>1% L-Glutamine<br>1% Penicillin-Streptomycin<br>10 ng/ml basic Fibroblast Growth Factor | ThermoFisher Scientific, 1634010<br>GE Healthcare, HYCLSH30070<br>ThermoFisher Scientific, 11140035<br>Sigma-Aldrich, M7522<br>ThermoFisher Scientific, 25030024<br>ThermoFisher Scientific, 15140122<br>Merck, GF003AF |
| Neural Induction Medium | Advanced DMEM/F12<br>1% N-2 Supplement<br>2 μg/ml Heparin<br>1% Penicillin-Streptomycin | ThermoFisher Scientific, 1634010<br>ThermoFisher Scientific, 17502048<br>Sigma-Aldrich, H3149<br>ThermoFisher Scientific, 15140122 |
| NPC Medium | Advanced DMEM/F12<br>1% N-2 Supplement<br>2% B-27 minus RA Supplement<br>1 μg/ml Laminin<br>1% Penicillin-Streptomycin<br>20 ng/ml basic Fibroblast Growth Factor | ThermoFisher Scientific, 1634010<br>ThermoFisher Scientific, 17502048<br>ThermoFisher Scientific, 12587010<br>Sigma-Aldrich, L2020<br>ThermoFisher Scientific, 15140122<br>Merck, GF003AF |
| Astrocyte Medium | Advanced DMEM/F12<br>1% N-2 Supplement<br>2% B-27 minus RA Supplement<br>1 μg/ml Laminin<br>1% Penicillin-Streptomycin<br>10 ng/ml Bone morphogenic protein 4<br>10 ng/ml Leukemia Inhibitory Factor | ThermoFisher Scientific, 1634010<br>ThermoFisher Scientific, 17502048<br>ThermoFisher Scientific, 12587010<br>Sigma-Aldrich, L2020<br>ThermoFisher Scientific, 15140122<br>BioVision, 4578<br>PreproTech, 300-05 |
| iCell Medium | BrainPhys Neuronal Medium<br>1% N-2 Supplement<br>1% iCell Nervous System Supplement<br>2% iCell Neural Supplement B<br>1 μg/ml Laminin<br>1% Penicillin-Streptomycin | Stemcell Technologies, 05790<br>ThermoFisher Scientific, 17502048<br>Cellular Dynamics, M1031<br>Cellular Dynamics, M1029<br>Sigma-Aldrich, L2020<br>ThermoFisher Scientific, 15140122 |
| *Ngn2* Medium | Neurobasal Medium<br>2% B-27 minus RA Supplement<br>1% Glutamax<br>10 ng/ml NT3<br>10 ng/ml BDNF<br>0.1 μg/ml primocin | ThermoFisher Scientific, 21103049<br>ThermoFisher Scientific, 12587012<br>ThermoFisher Scientific, 35050061<br>PeproTech, 450-03<br>ProSpec, CYT 207<br>Invivogen, ant-pm-1 |

**Figure 1:**
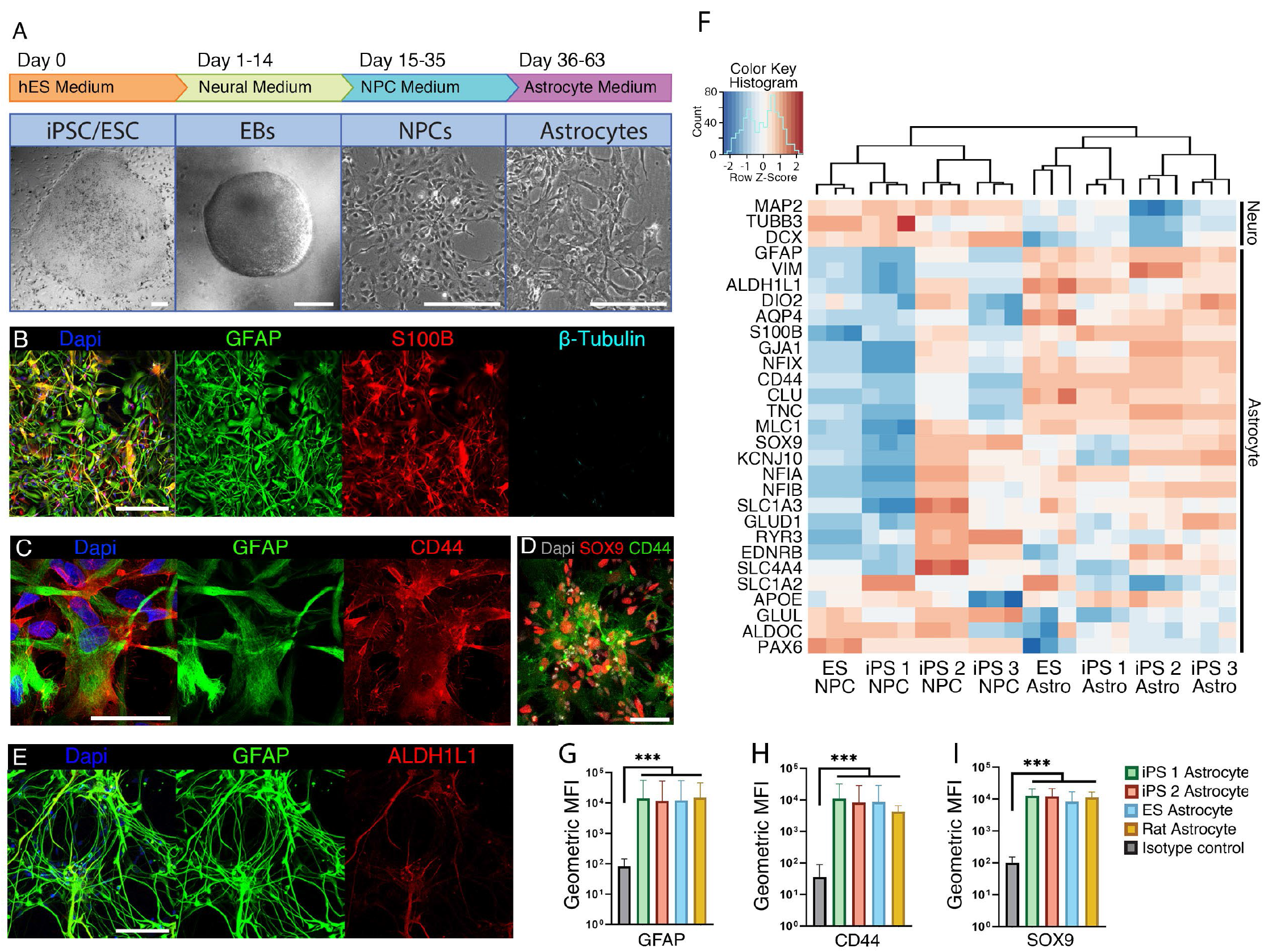
Astrocyte differentiation leads to a pure population of human astrocytes in four weeks. (**A**) Schematic representation of the differentiation protocol with representative DIC images of the different stages (scale bars = 200 μm). (**B**) Staining for astrocytic markers 4 weeks after NPC stage (GFAP, green and S100β, red) and an early neuronal marker (β-tubulin, cyan) (scale bar = 200 μm). (**C**) Staining for glial membrane marker CD44 (red) and GFAP (green) (scale bar = 50 μm). (**D**) Nuclear astrocytic marker SOX9 is shown in red, together with CD44 in green (scale bar = 50 μm). (**E**) A subset of astrocytes (GFAP, green) stains positive for ALDH1L1 (red) a mature astrocyte marker (scale bar = 100 μm). (**F**) Clustering of RNA sequencing data of 3 differentiation batches of NPCs to astrocytes from 4 different human pluripotent stem cell lines. Genes of interest are depicted on the left, on the top 3 neuronal genes and below 26 astrocyte genes. (**G**), (**H**), (**I**) Flow-cytometry data from iPS1, 2 and an embryonic stem cell (ES) line shows that 5-week-old astrocyte cultures show comparable geometric mean fluorescence intensity (MFI) to primary rat astrocytes.

### Immunocytochemistry

Cells were fixed using 4% formaldehyde (FA) in PBS (Merck, 1040032500) and labeled using immunocytochemistry. Primary antibody incubation was performed overnight at 4°C. Secondary antibodies were incubated for 2 hours at room temperature. Both primary and secondary antibody incubation were performed in staining buffer [0.05 M Tris, 0.9% NaCl, 0.25% gelatin, and 0.5% Triton X-100 (Sigma, T8787) in PBS (pH 7.4)]. Primary antibodies and their dilutions can be found in **Table 2**. Secondary antibodies conjugated to Alexa-488, Alexa-647 or Cy3 were used at a dilution of 1:400 (Jackson ImmunoResearch). Nuclei were visualised using DAPI (ThermoFisher Scientific, D1306). Samples were mounted using Mowiol 4-88 (Sigma-Aldrich, 81381) and imaged using a Zeiss LSM 800 confocal microscope (Oberkochen, Germany).

**Table 2:**
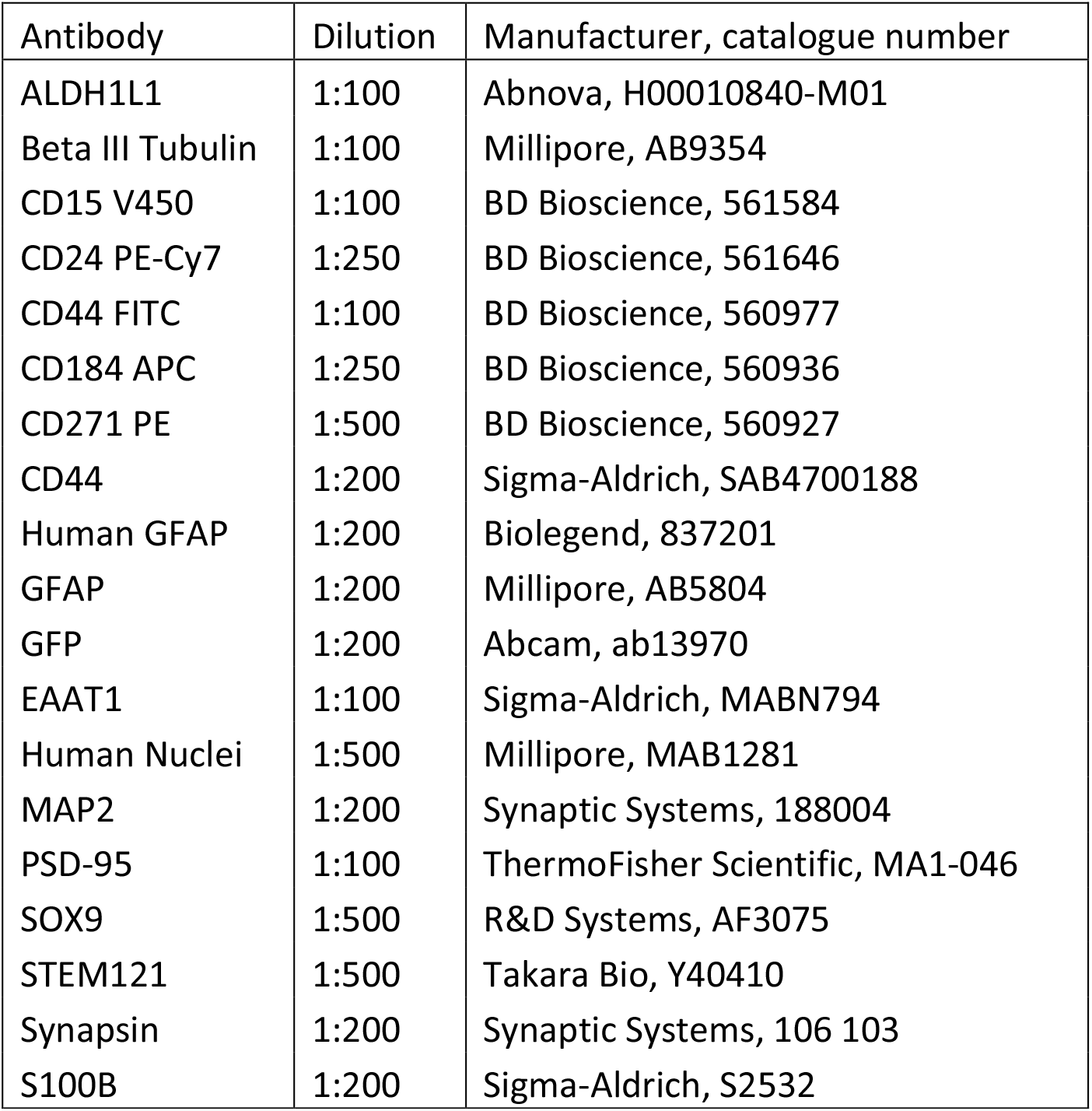
List of antibodies used.

### Flow cytometry quantification

Cells were detached from the culture dish using ACCUTASE™ (Stemcell technologies, 07920), washed, spun down, resuspended in PBS/2% FBS and stained with a primary antibody (**Table 2**) on ice for 30 minutes. Next, cells were washed and stained using a secondary antibody (Jackson ImmunoResearch, 1:400), kept on ice for 30 minutes, and washed 2 more times. Samples were analyzed on a LSRFortessa (BD Bioscience). Secondary antibody alone was used as an isotype control.

### RNA sequencing

Total RNA was isolated from NPCs and their derived astrocytes (4 lines, n=3 per cell type) using a RNeasy mini kit (Qiagen, 74104). RNA samples were prepped using TruSeq® Stranded mRNA Library kit (Illumina, 20020594). The resulting DNA libraries were sequenced according to the Illumina TruSeq Rapid v2 protocol on an Illumina HiSeq2500 sequencer. 50 bp reads were generated, trimmed and mapped against GRCh38 using HiSat2 (version 2.1.0), gene expression values were called using htseq-count (version 0.9.1). Sequencing resulted in at least 21.1M reads per sample, with at least 16.7M counts in the expression profile and 22.7-25.2k expressed genes. Analysis was performed using a custom R script.

### Culture dissociation and single-cell RNA sequencing

Three samples were used for single-cell RNA sequencing, 8-week-old astrocyte cultures, 7-day-old *Ngn2* neuronal cultures and 3-week-old astrocyte/neuron co-cultures. All derived from iPS1, for culture conditions, see Methods. For every sample, 4 cultures were pooled and dissociated using the Papain Dissociation System (Worthington Biochemicals, LK003150) according to manufacturer’s instructions. The single-cell solutions of iPSC-derived neural cultures were run on a Chromium Controller and final libraries were generated with the Chromium Next GEM Single cell 3’ reagents kits v3.1 (dual index) (PN-1000268, PN-1000120 PN-1000242, 10x Genomics) according to the manufacturer’s protocol. Libraries were sequenced on an Illumina Novaseq6000 system (28-10-10-90 cycles), aiming for 25000 reads/cell as suggested by 10X Genomics.

### Single-cell RNA sequencing data analysis

Sequenced samples were processed with the 10x Genomics Cellranger (v4.0.0) pipeline. There raw base call files were demultiplexed, reads were aligned (using STAR^24^, v2.5.1b) to human reference genome GrCH38 (v1.2.0) and filtered, finally barcodes and unique molecular identifiers were counted. Count data was processed using a custom pipeline developed with Seurat^25^ (v4.1.0) in the R statistical programming language (v4.0.5), code available on Github (https://github.com/kushnerlab/scRNAseqR) through request. Here we used the updated version of Seurat’s single cell transform^26^ (‘sctransform’) for normalization and variance stabilization. Integration was done by canonical correlation analysis on each mono- with co-culture. Highly variable genes were defined, these were used for dimension reduction through principal component analysis and Leiden algorithm^27^ for hierarchical clustering. Cell selection on cluster level was based on marker panels, for astrocytes: VIM, S100B and SOX9 and for neurons: MAP2, NEUROG2 and RBFOX3. Differential expression analysis was done with Seurat’s implementation of the Wilcoxon rank sum test. Clusters were annotated with SingleR^28^ (v1.4.1) by cross-referencing to a recently published database^29^, data was downloaded on June 9^th^, 2021. Pseudotime was calculated using Monocle3^30^ (v1.0.0).

### Astrocyte engraftment in neonatal Rag2^−/-^ mice

All mouse experiments were approved by the local animal welfare committee. IPSC-derived astrocytes were transplanted into immunodeficient neonatal (P1) *Rag2*^−/-^ BALB/c mice under cryoanesthesia. Roughly 5-10 × 10^4^ astrocytes were delivered in a 1 μl PBS-drop via a 1 mm-diameter pulled glass pipette into five different sites – posterior and anterior anlagen of the corpus callosum bilaterally and in the cerebellar peduncle dorsally^31^. Mice were sacrificed between 4 and 40 weeks of age by transcardiac perfusion with saline followed by 4% PFA. Brains were removed, left in 4% PFA for 2 hours at room temperature, transferred to a 10% phosphate buffer (PB 0.1 M, pH 7.3) and stored overnight at 4°C. Brains were embedded in 12% gelatin/10% sucrose blocks, fixation was performed for 2 hours at room temperature in a 10% pFA/30% sucrose solution. Embedded brains were stored, at least overnight, at 4°C before being sliced into 40 μm slices on a freezing microtome (Leica, Wetzlar, Germany; SM2000R). Brain sections were pre-incubated with a blocking buffer (0.5% Triton X-100 (Sigma, T8787) and 10% normal horse serum (NHS; ThermoFisher, 16050122) in PBS) for 1 hr at room temperature. Primary antibody incubation was done for 48 hours at 4°C. Secondary antibody incubation was performed for 2 hours at room temperature. Both primary and secondary antibody incubation were performed in a staining buffer (2% NHS and 0,5% Triton X-100 in PBS). Samples were mounted using Mowiol 4-88 (Sigma-Aldrich, 81381) and imaged using a Zeiss LSM 800 confocal microscope (Oberkochen, Germany).

### Human brain immunocytochemistry

All procedures with human tissue were performed with the approval of the Medical Ethical Committee of the Erasmus MC Rotterdam, including written consent of all subjects for brain donation in accordance with Dutch license procedures and the Declaration of Helsinki. Fresh-frozen post-mortem tissue blocks containing the middle frontal gyrus (BA9) from three donors (age 61-81) were obtained from the Erasmus MC Department of Pathology. Donors were confirmed to have no past medical history of any known psychiatric or neurologic illness, with additional confirmation of the absence of clinical neuropathology by autopsy examination^32^. Tissue blocks were postfixed for 7 days in 4% paraformaldehyde (0.1 M phosphate buffer, pH 7.3) at 4 °C. Tissue was subsequently transferred to 10% sucrose (0.1 M phosphate buffer, pH 7.3) and stored overnight at 4 °C. Embedding was performed in 12% gelatin/10% sucrose, with fixation in 10% paraformaldehyde/30% sucrose solution for 4 hours at room temperature and overnight immersion in 30% sucrose at 4 °C. Serial 40 μm sections were collected along the rostro-caudal axis using a freezing microtome (Leica, Wetzlar, Germany; SM2000R) and stored at −20 °C in a solution containing 37.5% ethylene glycol (Avantor, Central Valley, PA, USA, 9300), 37.5% glycerol (VWR Chemicals, Radnor, PA, USA, 24 386.298) and 25% 0.1 M phosphate buffer. Free-floating sections were washed thoroughly with PBS before being incubated in sodium citrate (10 mM) at 80 °C for 45 min and rinsed with PBS. Sections were pre-incubated with a blocking PBS buffer containing 1% Triton X-100 and 5% bovine serum albumin for 1 h at room temperature. Primary antibody labeling was performed in PBS buffer containing 1% Triton X-100 and 1% BSA for 72 h at 4 °C. Following primary antibody labeling, sections were washed with PBS and then incubated with corresponding Alexa-conjugated secondary antibodies and cyanine dyes (1:400, Braunschweig Chemicals, Amsterdam, The Netherlands) in PBS buffer containing 1% Triton X-100, 1% BSA for 4 h at room temperature. Nuclear staining was performed using DAPI (1:10 000, Thermo Fisher Scientific, Waltham, MA, USA). Images were acquired using a Zeiss LSM 800 confocal microscope (Oberkochen, Germany).

### Astrocyte size quantifications

Astrocytes were identified using a combination of antibodies (**Table 2**): (human) GFAP, STEM121 and human nucleus. Maximum projection images of 40 μm brain slices were used to determine cell size by drawing circles around typical protoplasmic astrocytes and calculating maximum diameter with Fiji (NIH ImageJ). Neighboring mouse astrocytes and transplanted human iPSC-derived astrocytes were analyzed in mice of different ages (4 - 40 weeks). Human astrocytes were analyzed in postmortem brain tissue (age 61-81).

### Glutamate uptake

Astrocytes (DIV28) were transduced with AAV9.GFAP.iGluSnFr.WPRE.SV40 (Penn Vector, AV-9-PV2914, 1×10^10^ vg/individual 24-wells). Two weeks after transduction, astrocytes were exposed to 50 μM glutamate (Sigma-Aldrich, G1251) in the culture medium, or co-cultured with iCell Glutaneurons (Cellular Dynamics, R1034) to measure their ability to take up glutamate. Fluorescence imaging was performed at 37°C/5% CO_2_ using a Zeiss LSM800 confocal microscope (Oberkochen, Germany) equipped with a live-imaging setup. Medium containing 50 μM glutamate was washed in after baseline recording. Co-cultures were recorded in iCell medium without addition of Glutamate. Fluorescence intensity was analyzed using Fiji (NIH ImageJ). ΔF/F was calculated for individual cells by manually drawing ROIs around fluorescent somata.

### Co-culture with iCell Glutaneurons

Human stem cell-derived astrocytes or rat astrocytes (Sciencell, SCCR1800) were grown in co-culture with iCell Glutaneurons (Cellular Dynamics, R1034). Astrocytes and neurons were plated in a 1:2 ratio on a 24-well Multiwell Electrode Array (MEA) plate (Multichannel Systems, 24W300-30G-288) or on coverslips, coated with Poly-L-Ornithine (Sigma-Aldrich, P4957) and 50 μg/ml laminin (Sigma-Aldrich, L2020). Co-cultures were maintained in iCell medium (**Table 1**) 37 °C/5%CO_2_ for up to 4 weeks.

### Co-culture with Ngn2-neurons in independent laboratory

iPSCs were directly differentiated into excitatory cortical layer 2/3 neurons by forcibly overexpressing the neuronal determinant *Neurogenin 2* (*Ngn2*)^14,33^. To support neuronal maturation, hPSC-derived astrocytes or freshly prepared rat astrocytes were added to the culture in a 1:1 ratio. At day 3, the medium was changed to *Ngn2*-medium (**Table 1**) and cytosine b-D-arabinofuranoside (Ara-C) (2 μM; Sigma, C1768) was added once to remove proliferating cells from the culture, facilitating long-term recordings of the cultures. From day 6 onwards, half of the medium was refreshed three times per week. Medium was additionally supplemented with 2.5% FBS (Sigma, F2442) to support astrocyte viability from day 10 onwards. Co-cultures were kept at 37 °C/5%CO_2_ throughout the entire differentiation process.

### Microelectrode Array (MEA) recordings

Spontaneous electrophysiological activity was recorded on a multiwell-MEA-system (Multichannel Systems). Plates were kept at 37°C and maintained in an air mixture containing 5% CO_2_. Plates were equilibrated to the chamber for 10 minutes and recorded for an additional 10 minutes. The signal was sampled at 10 kHz, filtered with a high-pass filter (i.e., Butterworth, 100 Hz cut-off frequency) and a low-pass filter (i.e. 4th order Butterworth, 3500 Hz cut-off frequency). The noise threshold was set at ±4.5 standard deviations. Recordings were analysed off-line using Multiwell-Analyzer (Multichannel Systems) and SPYCODE^34^.

### Electrophysiological recordings

Co-cultures were grown for 1-2 weeks, after which culture slides were transferred to the recording chamber and whole-cell patch-clamp recordings were performed as previously described^12^. Briefly, cultures were equilibrated to artificial cerebrospinal fluid (ACSF). In the recording chamber, slides were continuously perfused with ACSF at 1.5-2 mL/min, saturated with 95% O_2_/5% CO_2_ and maintained at 20-22°C. Recordings were performed with borosilicate glass recording micropipettes (3-6 MΩ). Data were acquired at 10 kHz using an Axon Multiclamp 700B amplifier (Molecular Devices), filtered at 3 kHz, and analyzed using pClamp 10.1 (Molecular Devices). Current-clamp recordings were performed at a holding potential of −70 mV. Intrinsic membrane properties were analysed using a series of hyperpolarizing and depolarizing square wave currents (500 msec duration, 1 sec interstimulus interval) in 5 pA steps, ranging from −30 to +30 pA. Data analysis was performed using a custom-designed script in Igor Pro-8.0 (Wavemetrics). Input resistance was calculated from the first two hyperpolarzing steps. Active properties were extracted from the first depolarizing step resulting in AP firing. AP threshold was defined by the moment at which the second derivative of the voltage exceeded the baseline. AP amplitude was measured from threshold. Neurons were categorized as “firing” if they were capable of firing 3 or more mature APs without significant accommodation during a depolarizing current step. Voltage-clamp recordings were performed at a holding potential of −80 mV. Synaptic events were detected using Mini Analysis Program (Synaptosoft, Decatur, GA). Bursting activity was defined as a period longer than 5 sec with a frequency of more than 20Hz, followed by a return to baseline.

### Synapse quantification

To quantify synapse formation, (co-)cultures were imaged 1 and 2 weeks after plating in different compositions: iCell Glutaneurons alone (1 week old, n=14), co-cultured with rodent astrocytes (1 week old, n=14, 2 weeks old, n=23) or co-cultured with iPSC-derived astrocytes (1 week old, n=35, 2 weeks old, n=24). Cultures were stained for synapsin, PSD95 and MAP2 (**Table 2**). Multiple (3-5) images were taken per coverslip. Synapses were counted as puncta with colocalization of PSD95, synapsin and MAP2 and normalized to MAP2 surface area per image (**Figure 5A**).

### Statistical analysis

Bulk RNA sequencing data was analyzed using the Fisher’s exact test, FDR corrected for multiple testing where necessary. For functional studies, statistical comparisons were performed using the Fisher’s exact test, a two-tailed t-test or analysis of variance (ANOVA), as indicated. Data are expressed as mean ± S.E.M., unless otherwise specified. The threshold for significance was set at *P*<0.05 for all statistical comparisons. Single cell RNA sequencing data was analyzed using Seurat’s implementation of Wilcoxon Rank Sum test. Here we set the parameters logfc.threshold and min.pct to 0. *P*-values adjusted for multiple testing error were used for thresholding significance at *P*<0.05.

## RESULTS

### Differentiation of human forebrain-patterned NPCs to astrocytes

Forebrain-patterned NPCs were generated from 3 iPSC-lines (iPS1-3) and an ESC-line as previously described^12^ with modifications. For all 4 lines, NPCs expressed SOX2 and Nestin, while MAP2 was rarely detected (**Supplementary Figure 1**). NPCs were cryopreserved and thawed in NPC medium at the start of a differentiation round. When confluent, cells were passaged 1:4 into astrocyte medium containing leukemia inhibitory factor (LIF) and bone morphogenetic protein 4 (BMP4) and grown for an additional 4 weeks (**Figure 1A, *Methods***). The resulting astrocytes expressed the canonical markers GFAP, S100B, SOX9, CD44, and were negative for neuronal markers such as β-tubulin and MAP2. A subset of astrocytes also expressed a marker for more mature astrocytes ALDH1L1 (**Figure 1B-E, Supplementary Figure 1**).

To further characterize the hPSC-derived astrocytes, and compare their expression profile to the parental NPCs, we performed bulk RNA sequencing of 3 independent differentiation batches of each of the 4 pluripotent stem cell lines (3 iPSC and 1 ESC). Hierarchical clustering based on Euclidean distance using variance stabilizing transformed (VST)-counts for established cell-type specific markers confirmed the distinction between NPC and astrocyte samples, and the batch-to-batch reproducibility (**Figure 1F**). Canonical astrocyte genes, e.g. *GFAP*, *AQP4*, and *S100B*, were robustly upregulated in astrocyte samples compared to NPC samples, while neuronal genes, e.g. *MAP2*, *TUBB3* and *DCX* were downregulated. Notably, astrocyte markers involved in the uptake and recycling of glutamate, e.g. *SLC1A3* and *GLUL*, were not robustly expressed in homogenous astrocyte cultures.

Flow cytometry based on GFAP, CD44 and SOX9 immunolabeling exhibited a similar geometric mean fluorescent intensity and cellular purity between hPSC-derived and primary rat astrocytes (**Figure 1G-I**, **Supplementary Figure 2**). These results together demonstrate that 4-week differentiation of NPCs in LIF / BMP4 is sufficient to obtain a relatively pure and homogenous population of hPSC-derived astrocytes.

### Single-cell RNA sequencing confirms maturation of astrocytes in neuronal co-culture

During brain development, astrocyte maturation is mutually co-regulated with neuronal maturation^35^. In order to investigate the transcriptomic impact of neuronal co-culture on hPSC-derived astrocytes, we performed single-cell RNA sequencing (scRNA seq) in three independent conditions: a) astrocyte mono-culture, b) neuronal mono-culture, and c) co-culture of astrocytes and neurons. As expected, the mono-culture and co-culture samples showed robust expression of their corresponding cell-type specific markers [e.g., *VIM* and *FABP7* (astrocytes); *MAP2* and *NEUROG2* (neurons)] (**Supplementary Figure 3A, B**).

We next calculated the Pearson correlation coefficient between the individual clusters in our samples and cell-type annotated clusters at different developmental stages (gestational week 14-25) in a recently published scRNAseq dataset of the developing human brain^29^. Developmentally, the transcriptional profile of both mono-cultures most closely resembled the cerebral cortex during gestational week 18 (**Supplementary Figure 3C, D**). Mono-culture hPSC-derived astrocytes included “radial glia” (38%), “dividing” (52%) and “endothelial” (10%) cells (**Supplementary Figure 3C, E**). Mono-culture neurons were most strongly correlated with excitatory neurons (90%), while the remaining 10% of cells had transcriptional profiles of intermediate progenitor cells (IPCs) (**Supplementary Figure 3D, F**). In the astrocyte-neuronal co-cultures, we observed 60% “excitatory neurons”, 20% “radial glia”, 13% “astrocytes” and 12% “dividing” cells (**Figure 2A, B**). Interestingly, the co-cultures yielded neurons with transcriptional profiles most strongly associated with gestational week 19 – slightly older than the mono-culture condition (**Figure 2A**). Neurons most closely resembled “excitatory neurons”, both in mono- and co-culture. In contrast, hPSC-derived astrocytes underwent a profound transcriptional adaptation in neuronal co-culture. Compared to astrocyte mono-culture, we observed fewer “dividing” cells, more “radial glia”, and a cluster observed exclusively in co-culture that was most strongly correlated with “astrocytes”.

**Figure 2:**
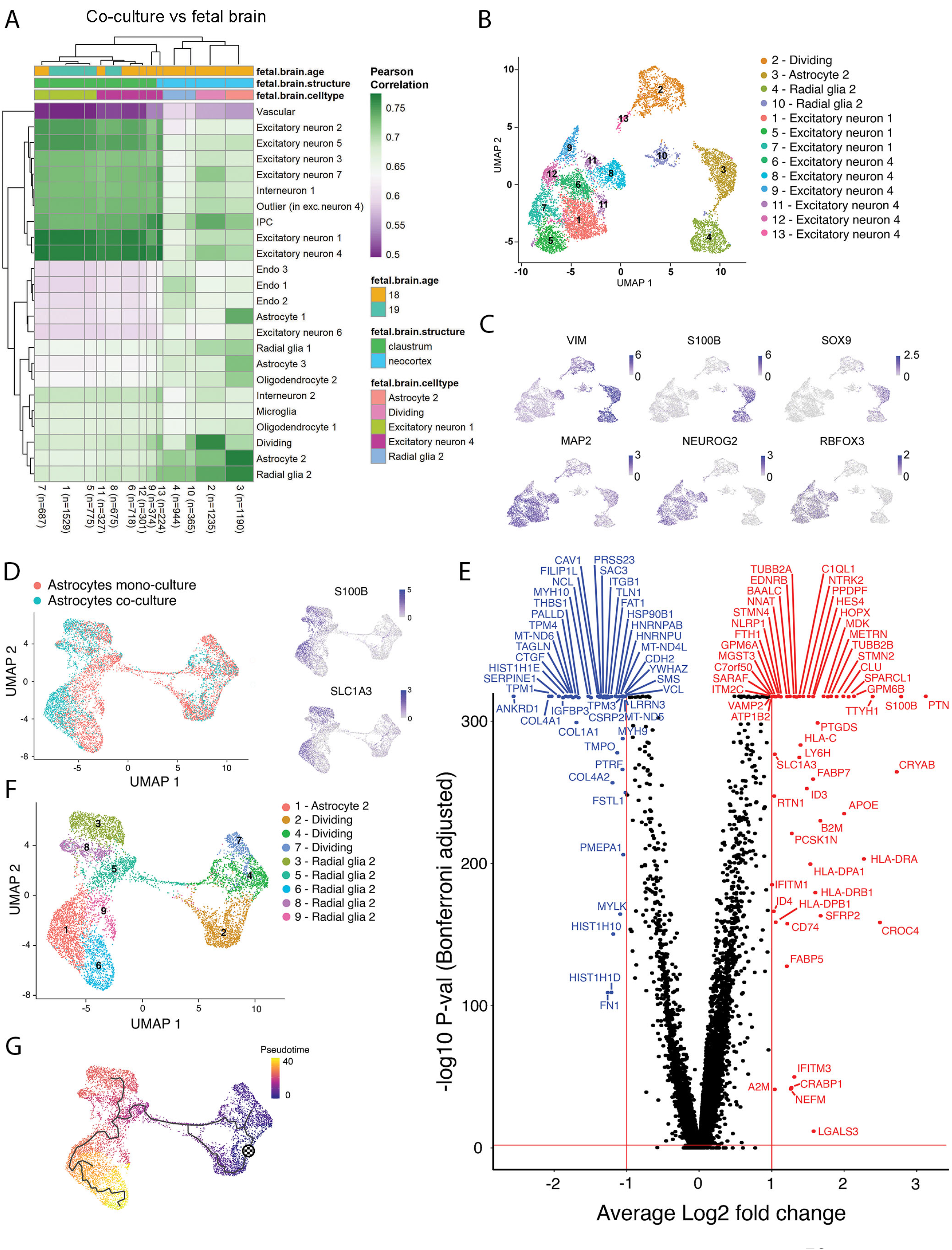
Single-cell RNA sequencing data confirms astrocyte identity and reveals transcriptional changes in co-culture conditions. (**A**) Heatmap showing Pearson’s correlation between scRNA seq cell-clusters of the co-culture sample and primary fetal brain tissue^29^. (**B**) UMAP projection of co-culture sample with transferred cell type labels with the highest correlation from primary brain tissue. (C) Expression of genes used for selection of astrocytes (top) and neurons from samples in the co-culture sample (bottom). (**D**) UMAP projection of integrated astrocyte sample with original sample identity indicated in green (co-culture) or red (mono-culture) (left). Expression of mature astrocyte markers *S100B* and *SLC1A3*, is skewed towards cells from the co-culture sample (right) (**E**) Volcano plot of significantly up- (red) and down-regulated (blue) genes in co-culture astrocytes vs mono-culture astrocytes. (**F**) Cell type label transfer on the UMAP projection of the integrated astrocyte sample. (**G**) Pseudotime trajectory of the integrated astrocyte sample suggests further maturation of astrocytes in the co-culture sample, pseudotime indicated from blue (early) to yellow (late).

In order to further characterize the maturation of hPSC-derived astrocytes, we next sought to identify genes that were differentially expressed in astrocytes from their corresponding mono-culture and co-culture conditions. We selected clusters in which >20% of cells expressed *VIM*, *S100B* and *SOX9* to create an integrated sample with mono- and co-culture astrocytes. Similarly, we used *RBFOX3*, *MAP2* and *NEUROG2* as markers to select neuronal clusters from mono-culture and co-culture conditions to generate a corresponding integrated neuron sample (**Figure 2C**). By retaining the original sample identity of individual cells in these integrated samples, we were able to directly compare their gene expression profiles (**Figure 2D, Supplementary Figure 3G**).

In the integrated astrocyte sample, some clusters showed a skewed distribution in the original sample identity. For example, clusters labelled “dividing” contained cells mainly from the mono-culture (mono-culture: 75.8%, co-culture: 24.2%), and in the cluster labelled “astrocyte” most cells originated from the co-culture sample (mono-culture: 16.8%, co-culture: 37.2%) (**Figure 2D, F, Supplementary Table 1**). No such skewing of clusters was observed in the integrated neuron sample (**Supplementary Figure 3C, Supplementary Table 2**).

Next, we performed differential expression analysis on the integrated samples to gain insight into the adaptations in astrocytes and neurons during co-culture (**Supplementary Tables 3** & **4**). **Figure 2E** displays a volcano plot of the differentially expressed genes of the integrated astrocyte sample with the most highly regulated genes shown in **Table 3**. As expected, many well-established markers of mature astrocytes were highly upregulated in hPSC-derived astrocytes from the co-culture sample, e.g. *S100B* and *SLC1A3* (**Figure 2D**). Moreover, multiple genes were found that are known to be specifically upregulated during astrocyte-neuron interactions and coordinated maturation, e.g. *SPARCL1*^36^, *METRN*^37,38^ and *CROC4*^39^.

**Table 3:**
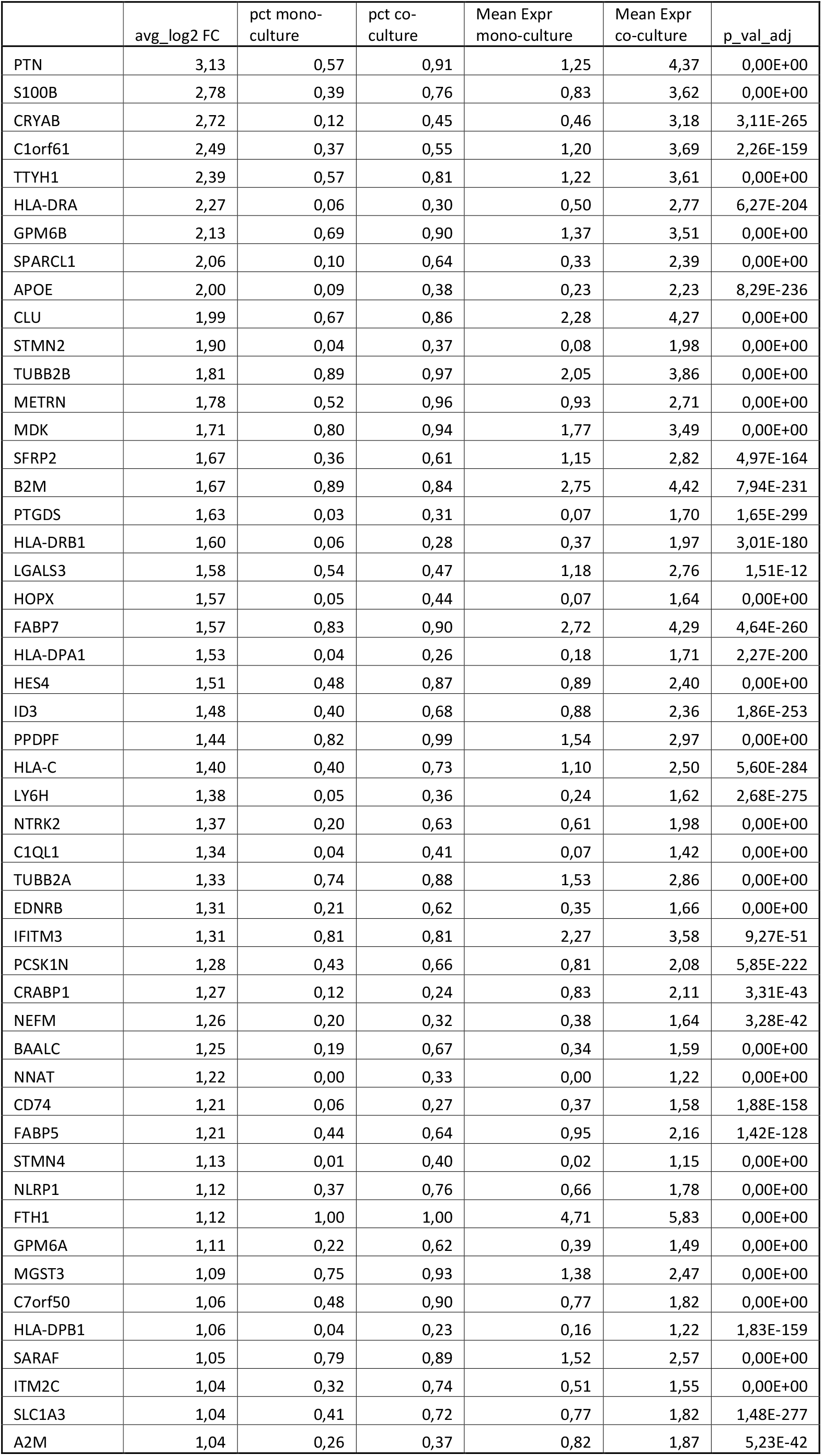
Top 50 differentially upregulated genes in co-culture astrocytes

Cross-referencing the differentially expressed genes from the integrated astrocyte sample with those of a study^8^ that investigated transcriptional differences between primary human astrocyte progenitors and mature astrocytes revealed a high correspondence (Fisher’s exact, *P*<0.001) (**Supplementary Table 5**), suggesting that astrocytes undergo additional maturation when grown with neurons. Pseudotime analysis of the integrated astrocyte sample confirmed this observation, in which “dividing” cells from the mono-culture sample are positioned at the beginning of the developmental trajectory and co-culture astrocytes enriched at the end (**Figure 2D, F, G**).

### hPSC-derived astrocytes retain a hominid morphology following transplantation into the mouse brain

Pluripotent stem cells have provided a unique opportunity to study the development and physiology of human brain cell lineages. This is especially valuable for astrocytes – the most highly dimorphic cell type between the higher order primates and other mammals^40^. Compared to their rodent counterpart, human astrocytes are larger and have a more complex morphology. We sought to investigate whether human astrocytes derived using our protocol maintain this feature when transplanted into the brains of neonatal immunodeficient *Rag2*^−/-^ mice. One week after transplantation, cells were mainly found near the subventricular zone (SVZ) of the lateral ventricles (**Supplementary Figure 4A)**. By 4 weeks after transplantation, cells had migrated away via the rostral migratory stream from the SVZ and could be found in the olfactory bulb (**Supplementary Figure 4B**). Eight weeks after transplantation, human cells were more globally distributed throughout the brain (**Figure 3A**). Eight months after transplantation, cells could still be found in the hippocampus and cortex (**Figure 3B**). Astrocytes *in vivo* inhabit non-overlapping anatomical and functional compartments, also known as astrocytic domains^7,41^. Transplanted hiPSC-derived astrocytes exhibited a similar cellular organization into astrocytic domains (**Figure 3A**). Moreover, we observed that transplanted astrocytes maintained their unique hominid features when transplanted into the mouse brain. The transplanted astrocytes had a larger maximum diameter (137.70 μm ± 10.3) (one-way ANOVA, *P*<0.001) compared to their rodent counterparts (29.28 μm ± 2.26), and similar (one-way ANOVA, p=0.99) to astrocytes (137.66 μm ± 5.48) in human post-mortem brain (**Figure 3C, D**).

**Figure 3:**
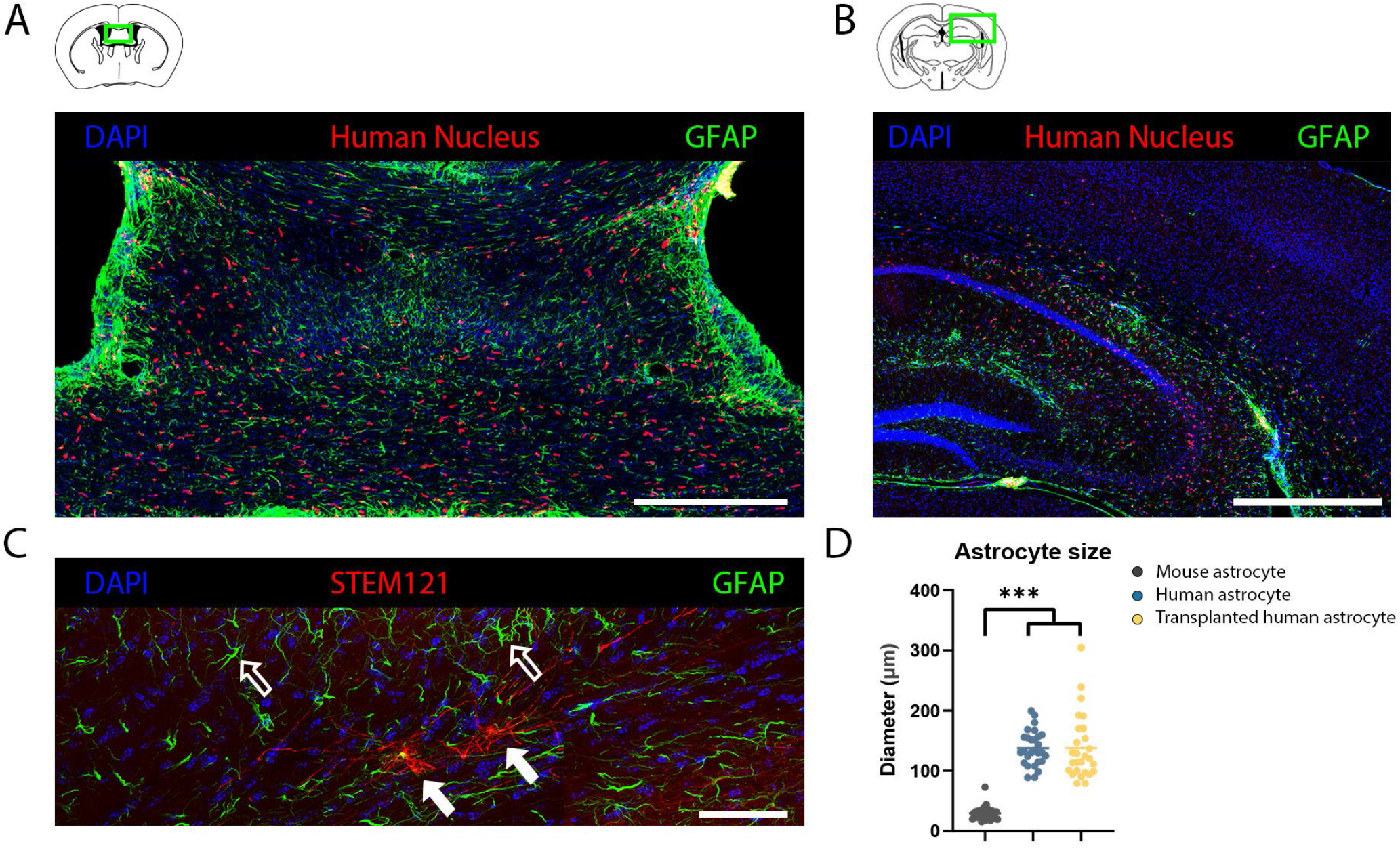
Human astrocytes integrate in the mouse brain after neonatal transplantation. (**A**) Human cells (red) arrange in astrocytic domains 8 weeks after transplantation (scale bar = 300 μm). (**B**) HPSC-derived astrocytes (red) 8 months after transplantation in the mouse hippocampus (scale bar = 500 μm). (**C**) Human astrocytes (red, solid arrows) are larger and more complex compared to their rodent counterpart (green, open arrows) in an identical *in vivo* environment (scale bar = 50 μm). (**D**) Cell size quantification (maximum diameter) of mouse astrocytes (black, n = 26), human astrocytes (blue, n = 28) and transplanted human astrocytes (yellow, n = 27).

### hPSC-derived astrocytes are able to buffer glutamate released in the synaptic cleft

To further characterize the functionality of the hPSC-derived astrocytes, we investigated their ability to buffer glutamate and support neuronal maturation. Glutamate uptake was assessed using iGluSnFr, a well characterized fluorescence-based intracellular glutamate sensor^42^. Astrocytes were transduced with an AAV9 vector expressing iGluSnFr under the control of the human GFAP promotor (**Figure 4A**, left panel). Astrocytes exposed to 50 μM extracellular glutamate exhibited a robust increase in fluorescence intensity (baseline: 0.9 × 10^3^ ± 3.2 au, glutamate: 2.0 ×10^3^ ± 2.0 au, *P*<0.01) (**Figure 4A-C**).

**Figure 4:**
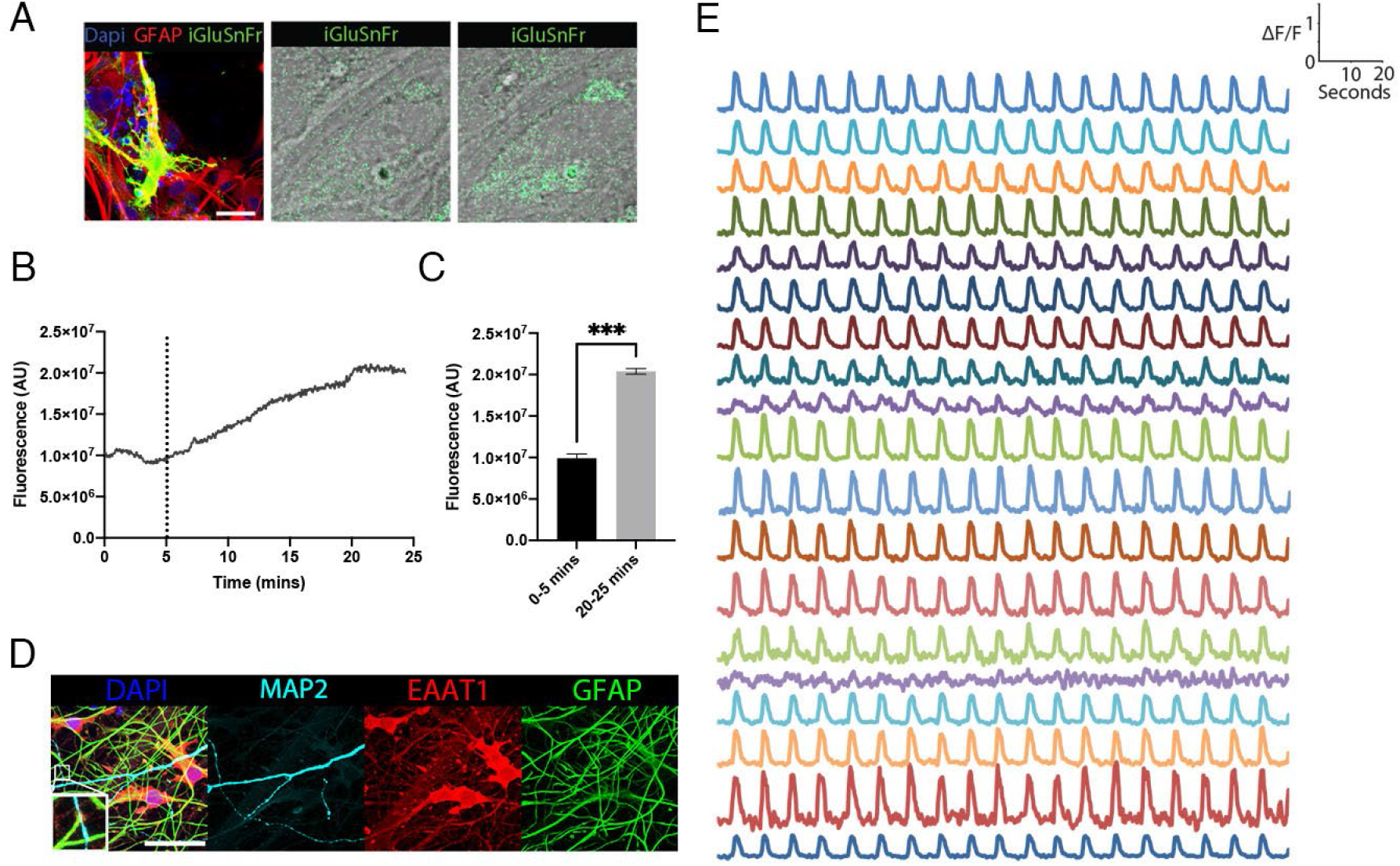
HPSC-derived astrocytes are able to buffer extracellular glutamate. (**A**) Glutamate sensor, iGluSnFr (green), expressed in astrocytes (red) (left panel) (scale bar = 25 μm), baseline fluorescence signal (middle panel), fluorescence signal after administration of 50 μM glutamate to the medium (right panel). (**B**) Time course of fluorescence signal over time, bath-application of 50 μM glutamate is indicated by the dashed line. (**C**) Comparison of the mean fluorescent signal of the first 5 minutes (0.9 × 10^3^ au ± 3.20) to the last 5 minutes of the recording (2.04 × 10^3^ au ± 2.04). (**D**) EAAT1 (red) expression increases in astrocytes (GFAP, green) when grown together with neurons (MAP2, cyan) (scale bar = 40 μm). (**E**) Traces of iGluSnFr fluorescent events of all individual cells in the field of view.

Astrocytes actively import extracellular glutamate through EAAT1 (*SLC1A3*)^43^. We found that EAATT1 expression was low when astrocytes are grown in mono-culture (**Supplementary Figure 5A**). In contrast, when co-cultured with neurons, EAAT1 (*SLC1A3*) expression increased two-fold (*P*<0.01) (**Figure 2D**). In addition, astrocytes seem to adopt a larger and more complex morphology when co-cultured with neurons (**Supplementary Figure 5B)**. Accordingly in co-culture, we also observed spontaneous repetitive synchronous increases of iGluSnFr fluorescence events (6.8 ± 0.36 min^−1^), a phenomenon that was not observed in mono-cultured astrocytes (**Figure 4D, Supplementary Movie 1**).

### hPSC-derived astrocytes promote the formation of functional synapses

Astrocytes are essential for proper neuronal network maturation and actively involved in the formation of synapses and establishment of network activity^44^. In order to investigate whether hPSC-derived astrocytes support neuronal network maturation, we quantified synapse formation and performed MEA and whole-cell patch-clamp recordings from mono-cultured neurons, co-cultured with primary rat astrocytes, or co-cultured with hPSC-derived astrocytes (**Figure 5, 6**).

**Figure 5:**
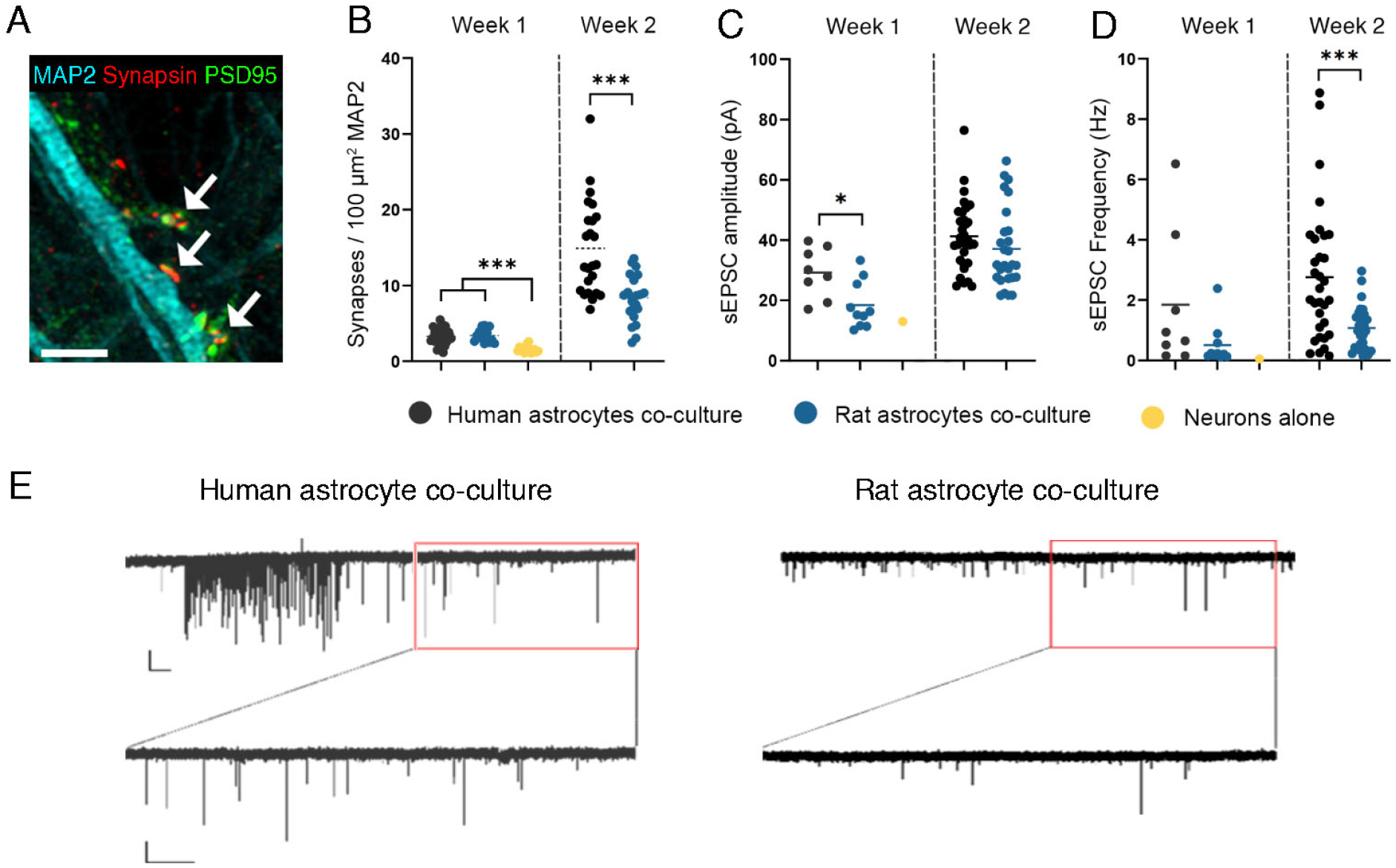
Synaptic networks are formed in co-cultures of neurons with astrocytes. (**A**) Representative image of synaptic staining used for quantification, co-localization of MAP2 (cyan), synapsin (red) and PSD95 (green) was counted as a synapse (arrows) (scalebar = 3 μm). (**B**) Synapse quantification of (co-)cultures at different timepoints using immunofluorescence. (**C, D**) Quantification of spontaneous EPSCs amplitude (**C**) and frequency (**D**) obtained using whole-cell patch-clamp recordings (week 1; n = 8 (human), n = 10 (rat) and n = 1 (neurons alone), week 2; n = 27 (human) and n = 32 (rat)). (**E**) Representative traces of a neuron from a co-culture with human (left) or rat (right) astrocytes, with a bursting event in the human co-culture, scale bar: 20 pA/5 sec.

**Figure 6:**
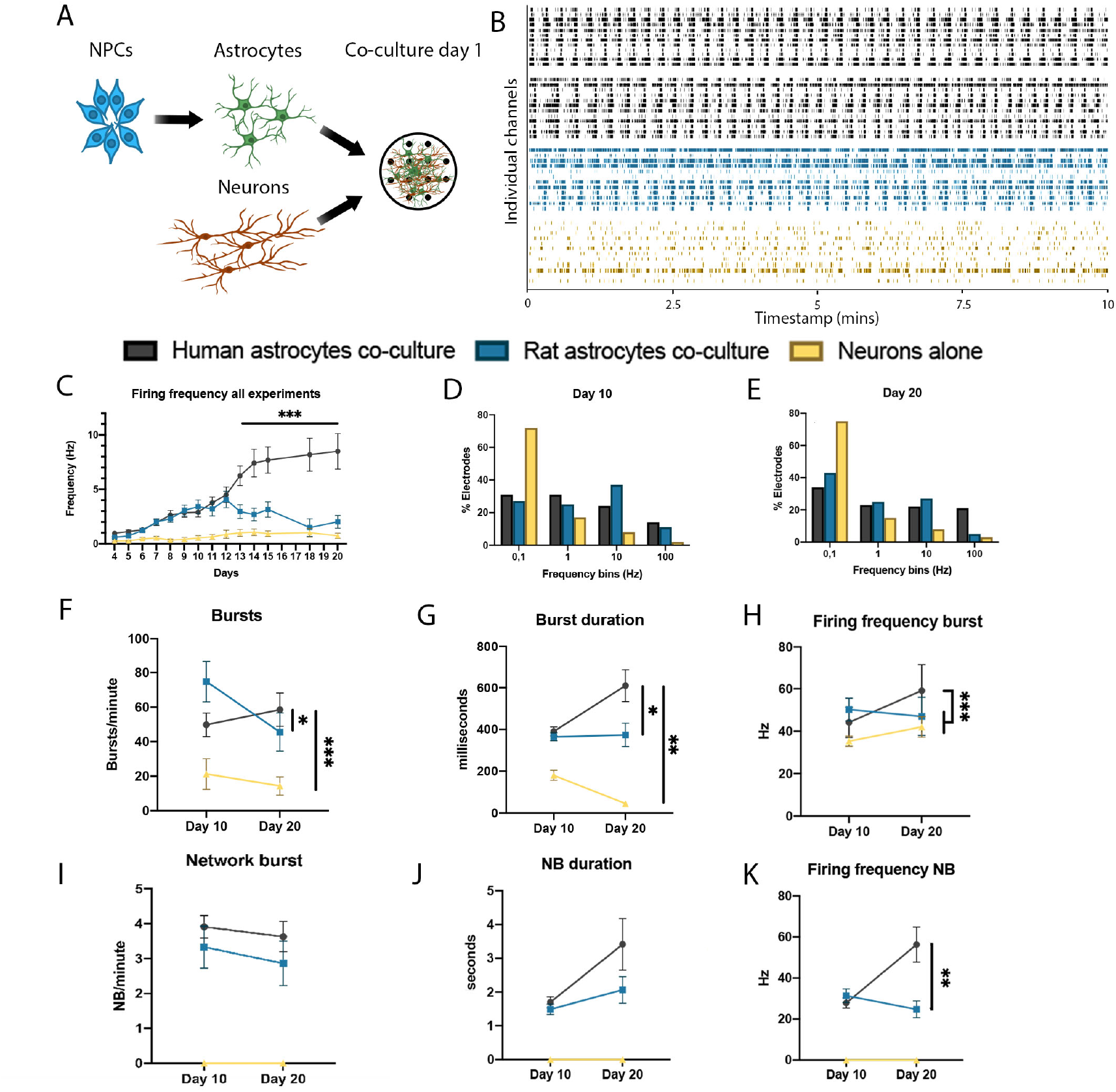
MEA recordings of (co-)cultures consisting of iPSC-derived neurons with or without astrocytes. (**A**) Experimental setup of neuronal co-cultures, NPCs (blue) are first differentiated to astrocytes (green) and plated together with neurons (red) in a co-culture. (**B**) Representative raster plots of spontaneous electrophysiological activity of individual wells in different co-culture conditions, with human astrocytes (black) from different batches, rodent astrocytes (blue) or neurons alone (yellow). (**C**) Mean firing frequency over time. (**D**), (**E**) Activity of all electrodes over multiple experiments presented in frequency bins 10 (D) and 20 (E) days after plating. **F, G, H** individual burst per minute (F), duration (G) and frequency (H). **I, J, K** Network burst per minute (I) duration (J) are similar between rodent and human astrocyte co-cultures, firing frequency (K) is increased in human astrocyte co-cultures.

One week after plating, we observed a greater number of synapses (**Figure 5A**) in co-cultures with hPCS-derived (3.3 ± 0.18/100 μm^2^ MAP2) or rat (3.4 ± 0.23/100 μm^2^ MAP2) astrocytes than in cultures with neurons alone (1.5 ± 0.12/100 μm^2^ MAP2) (*P*<0.001) (**Figure 5B**). At later timepoints, we were unable to keep neurons alive without supplementing astrocytes. After two weeks, co-cultures with hPSC-derived astrocytes had significantly more synapses (14.9 ± 1.27/100 μm^2^ MAP2) than those with rat astrocytes (8.39 ± 0.63/100 μm^2^ MAP2) (*P*<0.001).

Using whole-cell patch-clamp recordings, we found robust spontaneous excitatory post-synaptic currents (sEPSCs) in cultures supplemented with astrocytes, while EPSCs were nearly undetectable in cultures without astrocytes. One week after plating, sEPSC amplitude was significantly higher in co-cultures with hPSC-derived astrocytes (29.21 pA ± 2.75) compared to rat astrocytes (18.42 pA ± 2.37) (*P*<0.05) (**Figure 5C**). By two weeks after plating, sEPSC amplitude showed a non-significant increase in co-cultures with hPSC-derived (41.24 pA ± 1.97) versus rat (37.05 pA ± 2.51) astrocytes (*P* = 0.19) (**Figure 5C**). In one week old cultures, sEPSC frequency showed a non-significant increase in co-cultures with hPSC-derived (1.85 Hz ± 0.81) versus rat (0.51 Hz ± 0.21) astrocytes (*P* = 0.09) (**Figure 5D**). By two weeks after plating, sEPSC frequency was significantly increased in both conditions compared to week 1 (*P*<0.001). Moreover, EPSC frequency was significantly increased in co-cultures with hPSC-derived astrocytes (2.76 ± 0.39 Hz) compared to those with rat astrocytes (1.07 ± 0.14 Hz) (*P*<0.001) (**Figure 5D**). We found a non-significant increase in neurons exhibiting sEPSC burst activity in co-cultures with hPSC-derived astrocytes (38.24%) versus rat astrocytes (20.69%) (Fisher’s exact, *P*=0.17) (**Supplementary Figure 6G, H**). EPSC ride time was slower in co-cultures with hPSC-derived astrocytes (1.76 ms ± 0.08) versus rat astrocytes (1.46 ms ± 0.09) (*P*<0.05) (**Supplementary Figure 6P**) We observed no statistically significant differences in the intrinsic properties of neurons in either of the co-culture conditions (**Supplementary Figure 6**).

### Formation of high-frequency network activity in hPCS-derived astrocyte and neuron co-cultures

The developmental time course of network activity was further studied using MEA recordings (**Figure 6A)**. Raster plots showing representative activity for individual wells can be seen in **Figure 6B**. Neurons co-cultured with astrocytes exhibited increased activity compared to neuronal mono-cultures, beginning 5 days after plating (**Figure 6C**). This difference became even more pronounced over time, reaching a plateau after 2 weeks of activity. Notably, the firing rate was higher in co-cultures with hPSC-derived versus primary rat astrocytes, beginning 12 days after establishment of the cultures (2-way ANOVA, *P*<0.005). The firing frequency of all individual electrodes was binned across 10 days (**Figure 6D**) or 20 days (**Figure 6E**) after plating the cultures on MEAs (**Figure 6D, E**). Twenty days after plating, 21% of the electrodes in hPSC-derived astrocyte co-cultures exhibited a firing rate >100 Hz. In contrast, for co-cultures with rat astrocytes, only 5% of the electrodes recorded firing frequencies over 100 Hz (Fisher’s exact, *P*<0.01). In addition to overall firing rates, we also quantified individual burst activity (**Figure 6F-H**). At day 20, bursts occurred more frequently in neuronal co-cultures with hPSC-derived astrocytes (58.56 bursts/min ± 9.66) compared to rat co-cultures (45.56 bursts/min ± 11.12) or without astrocytes (14.33 bursts/min ± 5.30) (**Figure 6F**). Bursts were also of a longer mean duration in neuronal co-cultures with hPSC-derived astrocytes (609.93 ms ± 76.00) compared to rat astrocytes (373.94 ms ± 55.57) or without astrocytes (44.84 ms ± 7.29) [*P*<0.05 (rat), *P*<0.01 (without astrocytes)] (**Figure 6G**), and exhibited a higher within-burst firing frequency [hPSC-derived (59.13 Hz ± 2.20), rat (46.98 Hz ± 2.32), without astrocytes (42.01 Hz ± 1.97); *P*<0.001 (rat), *P*<0.005 (without astrocytes)] (**Figure 6H**).

Network bursts (NBs) were defined as simultaneous bursts in at least 50% of all electrodes in a well. NBs were only detected in co-cultures, with no detectable NBs in neuronal cultures without astrocytes. At day 20, co-cultures using hPSC-derived and rat astrocytes NBs were similar in frequency (3.63 ± 0.44 min^−1^ (hPSC-derived), 2.86 ± 0.64 min^−1^ (rat); (*P*=0.19) **Figure 6I**) and duration (3.4 ± 0.77 sec (hPSC-derived), 2.1 ± 0.39 sec (rat); (*P*=0.10) **Figure 6J**). However, firing frequency within NBs was higher in co-cultures with hPSC-derived astrocytes (56.31 ± 8.56 Hz) compared to rat (24.77 ± 4.04 Hz) (*P*<0.01; **Figure 6K**).

In an effort to evaluate the robustness of the protocol, an independent laboratory (Nijmegen) compared their standardized workflow for MEA-based recordings of co-cultures of rat astrocytes and human *Ngn2*-induced neurons by substituting with hPSC-derived astrocytes (**Supplementary Figure 7A**)^45–47^. Neuronal firing frequency was significantly increased in co-cultures with hPSC-derived versus rodent astrocytes (2-way ANOVA, *P*<0.001) (**Supplementary Figure 7B**). Furthermore, human astrocyte co-cultures showed an increased NB frequency (*P*<0.005) (**Supplementary Figure 7C**) and similar NB duration (*P*=0.70; **Supplementary Figure 7D**).

## DISCUSSION

Here, we demonstrate a novel method to establish functional hPSC-derived astrocytes. Immunocytochemistry and bulk RNA sequencing reveals a relatively homogenous population of cells with widespread expression of canonical astrocyte markers such as GFAP, S100B, SOX9 and CD44 (**Figure 1**). Single-cell RNA sequencing shows that the transcriptional profile of these cells is most similar to primary human “radial glia”, “astrocytes” and a group of “dividing” cells (**Figure 2A, B, Supplementary Figure 3A, C**). Following transplantation in the mouse brain, hPCS-derived astrocytes maintain their uniquely hominid^7^ morphological characteristics (**Figure 3D**). Functional studies demonstrate that this protocol yields hPCS-derived astrocytes that are able to buffer excess glutamate from the synaptic cleft (**Figure 4E**), support the formation of functional synapses (**Figure 5**) and the establishment of active neural networks (**Figure 6**).

Astrocytes are essential for neuronal network development, survival and electrophysiological maturation^48,49,50^. Accordingly, there is a widely acknowledged pressing need for protocols to obtain astrocytes from human pluripotent stem cells, demonstrated by the many protocols that currently exist^16,17,18,19,15,51^. Two recent studies^52,53^ have demonstrated that supplementing additional iPSC-derived astrocytes to a neural culture containing neurons, astrocytes and NPCs improves the electrophysiological properties of neurons. Protocols to differentiate iPSCs to a homogenous population of astrocytes that support neuronal maturation can be time-consuming and challenging. The protocol reported here demonstrates the ability to rapidly and efficiently differentiate human (induced) pluripotent stem cells into functional astrocytes. By making use of an intermediate stage of NPCs that can be expanded and survives cryopreservation, astrocyte-cultures can be established while maintaining the genomic integrity of the iPSC-line.

Compared to their rodent counterpart, human astrocytes are larger and have a more complex morphology. A specific subtype of astrocyte, interlaminar astrocytes, are found exclusively in higher order primates^54^. It has been established that primary human astrocytes also functionally show differences with rodent astrocytes^8,9^. We demonstrate that astrocytes differentiated using our protocol maintain these morphological hominid characteristics. When transplanted into a rodent brain, they are larger and more complex compared to neighbouring mouse astrocytes. Interestingly, we observed a change in *in vitro* morphology of astrocytes becoming larger and more complex when grown in a co-culture with neurons combined with a profound change in their expression profile (**Figure 2, Supplementary Figure 4**). Together, these findings demonstrate a critical bidirectional interaction between neurons and astrocytes.

Even though we observe a relatively uniform expression of GFAP, S100B and SOX9 in pure hPSC-derived astrocytes, the scRNAseq data suggests distinct glial subtypes exist within this *in vitro* population, especially in co-culture with neurons. IPSC-derived astrocytes seem to undergo an additional maturation step when co-cultured with neurons and can be separated into distinct clusters (**Figure 2**). *In vivo*, astrocytes are divided into subtypes based on their morphology^40^. Here we show that it is also possible to make a distinction *in vitro* based on their transcriptome profile. Notably, in the co-culture sample, cluster 10 (“radial glia”) showed a strong increase in *APOE* expression (log2fold=4.8, *P*<0.01) compared to all other clusters (**Supplementary Figure 3H**), in the astrocyte mono-culture we did not observe a defined cluster with *APOE* expression (**Supplementary Figure 3I**). The identification of such a specific cluster of cells provides future opportunities for studies focused on the aetiology of Alzheimer’s disease and highlights the importance of establishing adequate culture conditions when using iPSC-derived cultures to study genes of interest.

One of the major characteristics of astrocytes is their ability to buffer glutamate from the synaptic cleft^55^ and consequently ensure proper signalling via the tripartite synapse. We detected *EAAT1* expression in our RNA sequencing data and observed robust EAAT1 staining in co-cultured astrocytes. Functionally, we observed iGluSnFr fluorescence events (~6.8 min^−1^) in a similar order of magnitude as the timing of network bursts recorded during MEA experiments (~3.7 min^−1^; **Figures 4E & 6I**), strongly suggesting that endogenous glutamate release by neurons is taken up by neighboring astrocytes with high temporal precision.

A widely adopted protocol^14^ that quickly yields a pure population of neurons was predicated on the addition of primary rodent astrocytes for the neurons to mature enough to perform electrophysiological measurements^56^. Here, we demonstrate that with our protocol it becomes possible to create a fully functional human PSC-derived neural co-culture system. This provides the opportunity to precisely control the culture composition of a fully human population of neural cells, making it possible to study the effects of cell-type specific genotypes or targeted genetic manipulations of co-cultured astrocytes and/or neurons. In addition, we show that astrocytes generated using our protocol can be efficiently integrated into the existing workflow of an independent laboratory, emphasizing the robustness of the method and encouraging its broad adoption in the field.

Our experiments show that iPSC-derived neurons are more active and receive more synaptic input in a co-culture with human versus rodent astrocytes. Synapse formation is accelerated in a fully human co-culture system, resulting in an increased detection of sEPSCs through whole-cell electrophysiology (**Figure 5**). On a MEA, we observe a higher firing frequency in co-cultures with hPSC-derived astrocytes compared to primary rat astrocytes, both for individual events and within (network) bursts (**Figure 6**). This increased firing frequency can be explained by a subset (21%) of electrodes that recorded a firing frequency >100 Hz in the hPCS-derived astrocyte co-cultures, compared to 5% of electrodes with a similar firing frequency in the rat astrocyte co-cultures. We were unable to detect any substantial synapse formation or neuronal activity in cultures without astrocytes and had great difficulty maintaining these cultures for prolonged periods of time, illustrating the crucial role astrocytes play in neuronal maturation and network formation.

Taken together, our findings demonstrate that using this protocol we are able to create functional hPCS-derived astrocytes from NPCs in 28 days. The protocol presented here provides an opportunity to efficiently derive functional astrocytes from human pluripotent stem cells that compare favourably to primary rodent astrocytes.

## Acknowledgements

This work was supported by the Netherlands Organ-on-Chip Initiative, an NWO Gravitation project (024.003.001) funded by the Ministry of Education, Culture and Science of the government of the Netherlands (S.A.K., F.M.S.D.V., B.L.), ERA PerMed Joint Transnational Call – ZonMw 456.008.003 to S.A.K., an Erasmus MC Human Disease Model Award to F.M.S.D.V., and by a Simons Foundation grant (SFARI) #890042 to N.N.K.

Authors declare no conflicts of interest

