## Supplementary figures + tables for "Rapid specification of human pluripotent stem cells to functional astrocytes"

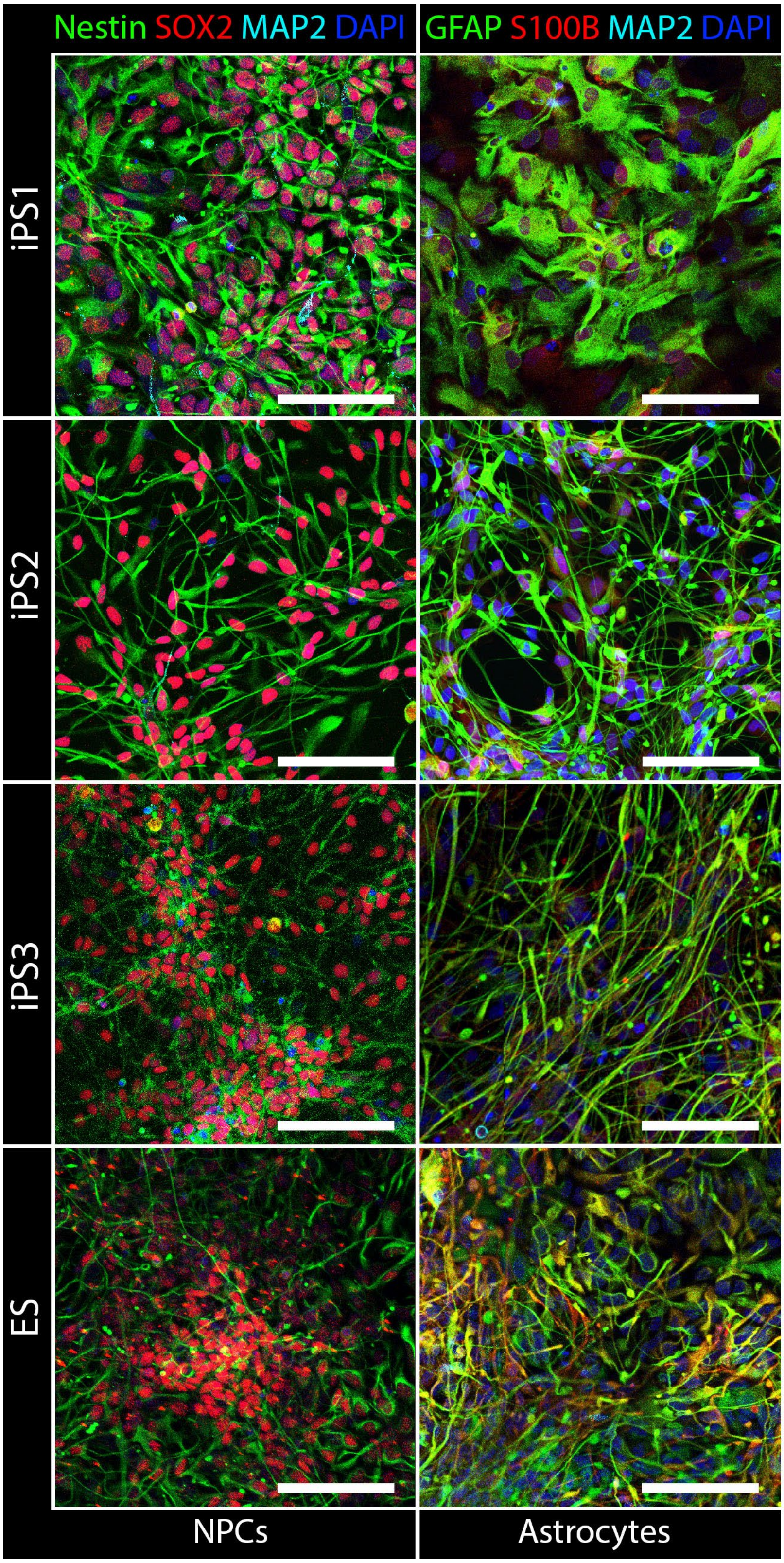


**Supplementary Figure 1: Immunofluorescent labeling of NPCs and their derived astrocyte cultures.** NPCs stain positive for Nestin (green) and SOX2 (red) and negative for MAP2 (cyan). Astrocytes stain positive for GFAP (green) and S100B (red) and negative for MAP2 (cyan) (scale bar = 50 μm).

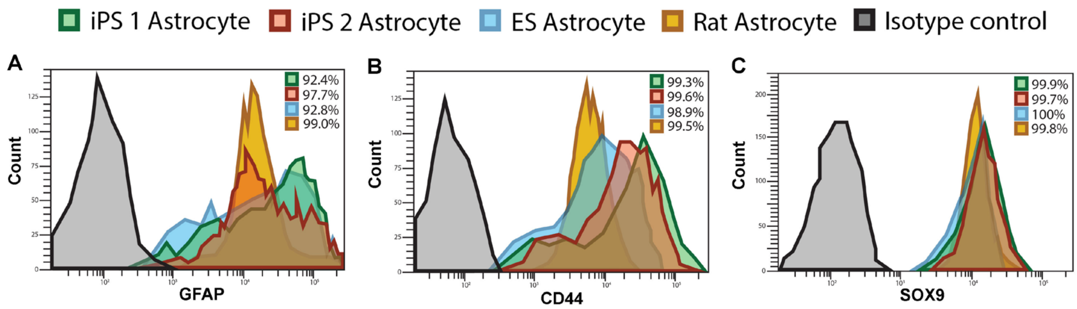


**Supplementary Figure 2: Flow-cytometry quantification of astrocyte markers.** Fluorescence intensity histogram plots for iPS 1-, iPS 2- and embryonic stem cell (ES)-derived astrocytes compared to primary rat astrocytes for GFAP (**A**), CD44 (**B**) and SOX9 (**C**).

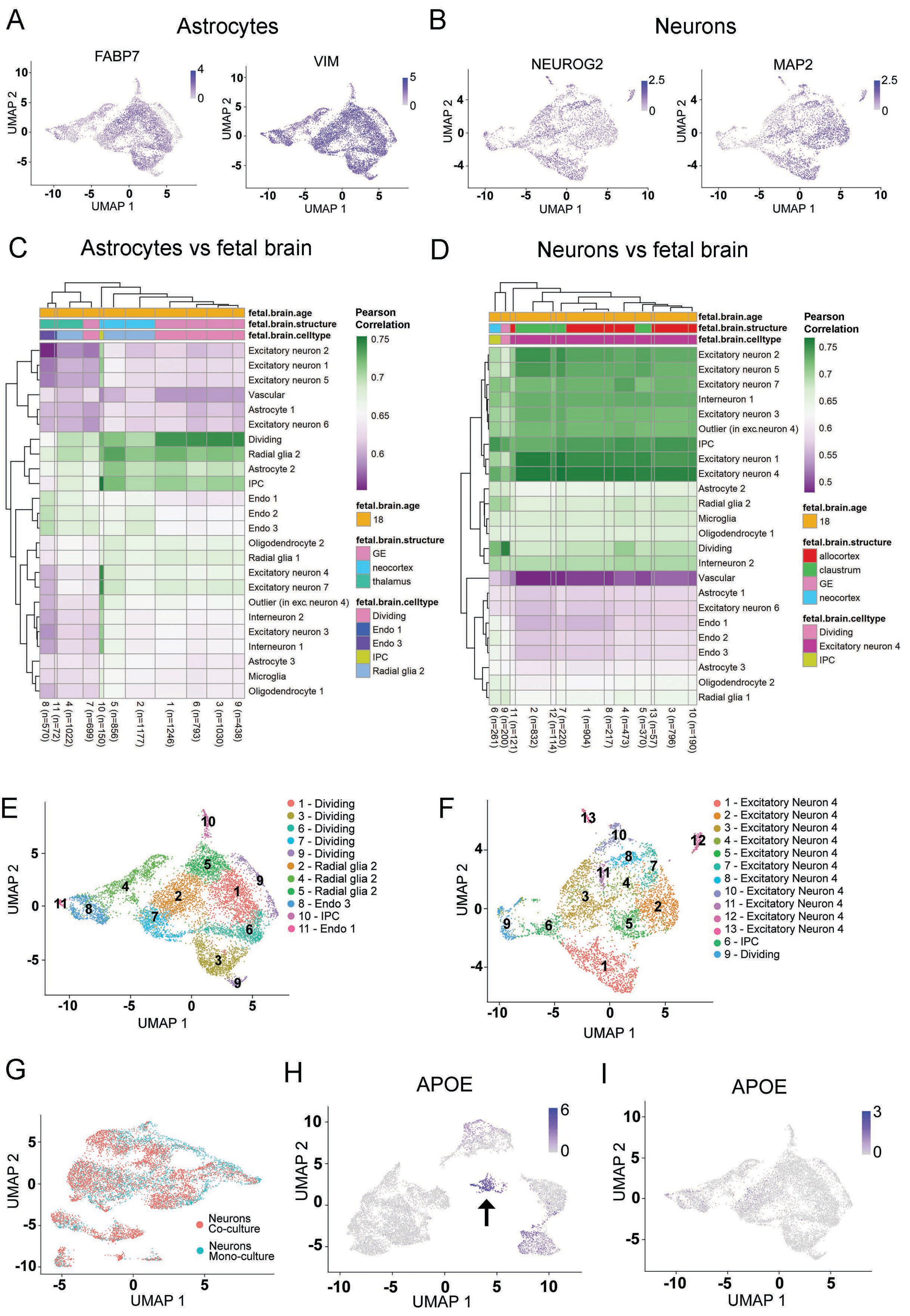
A
Astrocytes
FABP7
VIM
B
Neurons
NEUROG2
MAP2
C
Astrocytes vs fetal brain
D
Neurons vs fetal brain
fetal.brain.age
fetal.brain.structure
fetal.brain.celltype
Pearson Correlation
GE
neocortex
thalamus
allocortex
claustrum
Dividing
Endo 1
Endo 3
IPC
Radial glia 2
Excitatory neuron 4
E
1 - Dividing
3 - Dividing
6 - Dividing
7 - Dividing
9 - Dividing
2 - Radial glia 2
4 - Radial glia 2
5 - Radial glia 2
8 - Endo 3
10 - IPC
11 - Endo 1
F
1 - Excitatory Neuron 4
2 - Excitatory Neuron 4
3 - Excitatory Neuron 4
4 - Excitatory Neuron 4
5 - Excitatory Neuron 4
7 - Excitatory Neuron 4
8 - Excitatory Neuron 4
10 - Excitatory Neuron 4
11 - Excitatory Neuron 4
12 - Excitatory Neuron 4
13 - Excitatory Neuron 4
6 - IPC
9 - Dividing
G
Neurons Co-culture
Neurons Mono-culture
H
APOE
I
APOE
UMAP 1
UMAP 2

**Supplementary Figure 3: Single-cell RNA sequencing data from astrocyte and neuron mono-culture samples.** (**A**) Astrocyte sample shows homogenous expression of known astrocyte markers, e.g. FABP7 and VIM. (**B**) *Ngn2*-neuron sample shows homogenous expression of known neuronal markers, e.g. MAP2 and NEUROG2. (**C**) Heatmap showing Pearson's correlation between scRNA seq cell-clusters of the astrocyte sample and primary fetal brain tissue. (**D**) Heatmap showing Pearson's correlation between scRNA seq cell-clusters of the *Ngn2*-neuron sample and primary fetal brain tissue. (**E**) UMAP projection of the astrocyte sample with transferred cell type labels with the highest correlation from primary brain tissue. (**F**) UMAP projection of the *Ngn2*-neuron sample with transferred cell type labels with the highest correlation from primary brain tissue. (**G**) UMAP projection of integrated *Ngn2*-neuron sample with original sample identity indicated in green (mono-culture) or red (co-culture). (**H, I**) APOE expression is upregulated in astrocytes under co-culture conditions and mostly expressed in a single cluster (cluster 10, "Radial glia", arrow) (**H**), in a culture with only astrocytes (**I**) APOE expression is lower.

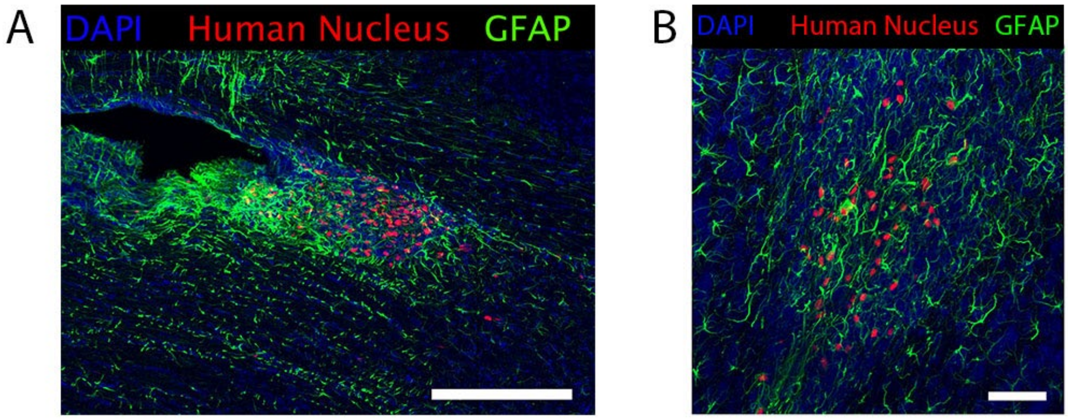


**Supplementary Figure 4: IPSC-derived astrocytes transplanted in a mouse brain.** (**A**) 4 weeks after transplantation human astrocytes are mainly found in the subventricular zone of the lateral ventricles (scale bar = 200 μm). (**B**) Human astrocytes in the olfactory bulb of a 4-week-old mouse (scale bar = 50 μm).

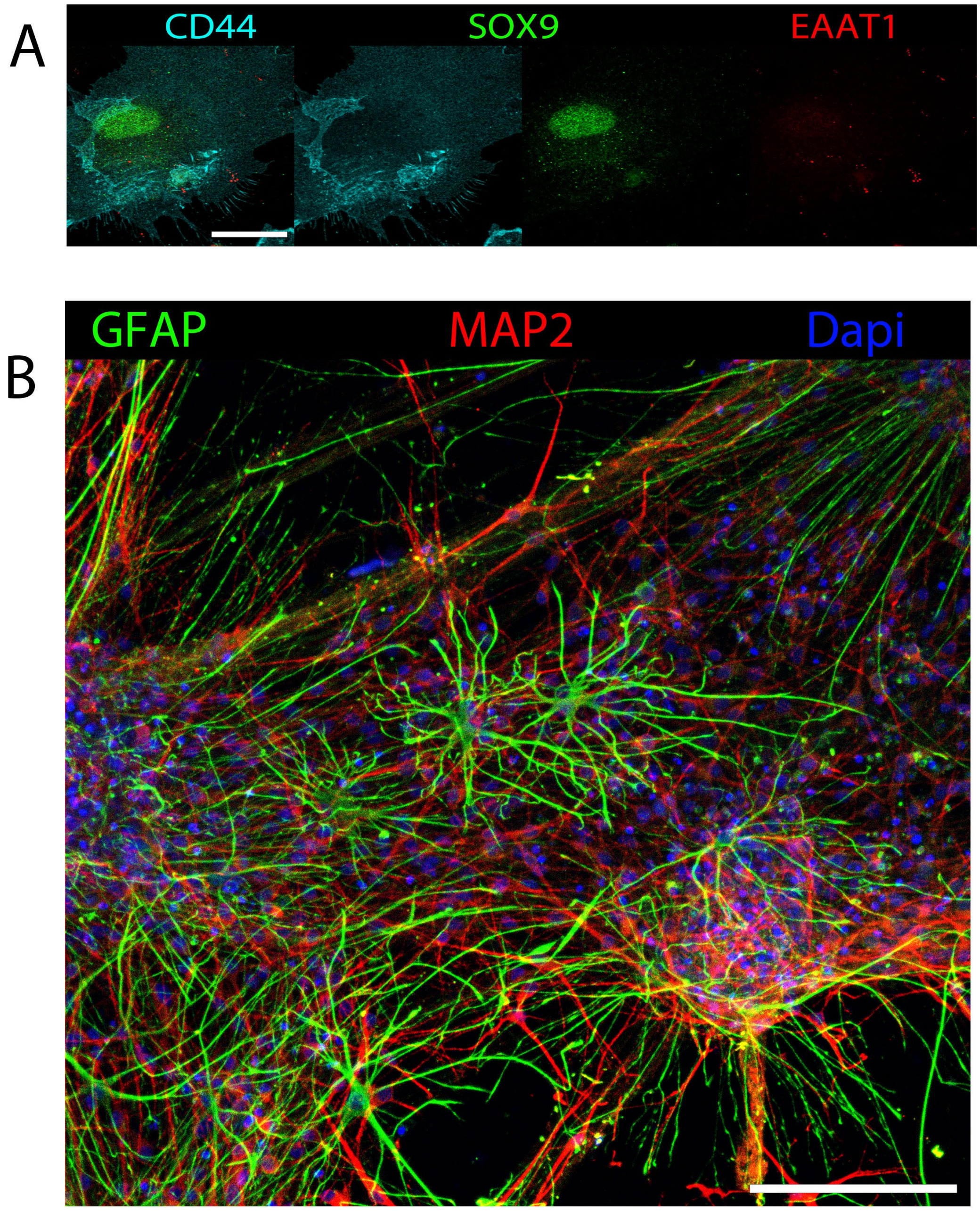


**Supplementary Figure 5: Astrocyte EAAT1 expression and morphology changes upon co-culture with neurons.** (**A**) 11-week-old astrocyte in mono-culture shows little EAAT1 staining (scale bar = 20 μm). (**B**) Astrocytes become larger and more branched when grown in a co-culture compared to a pure population (**Figure 1B-E**) (scale bar = 100 μm).

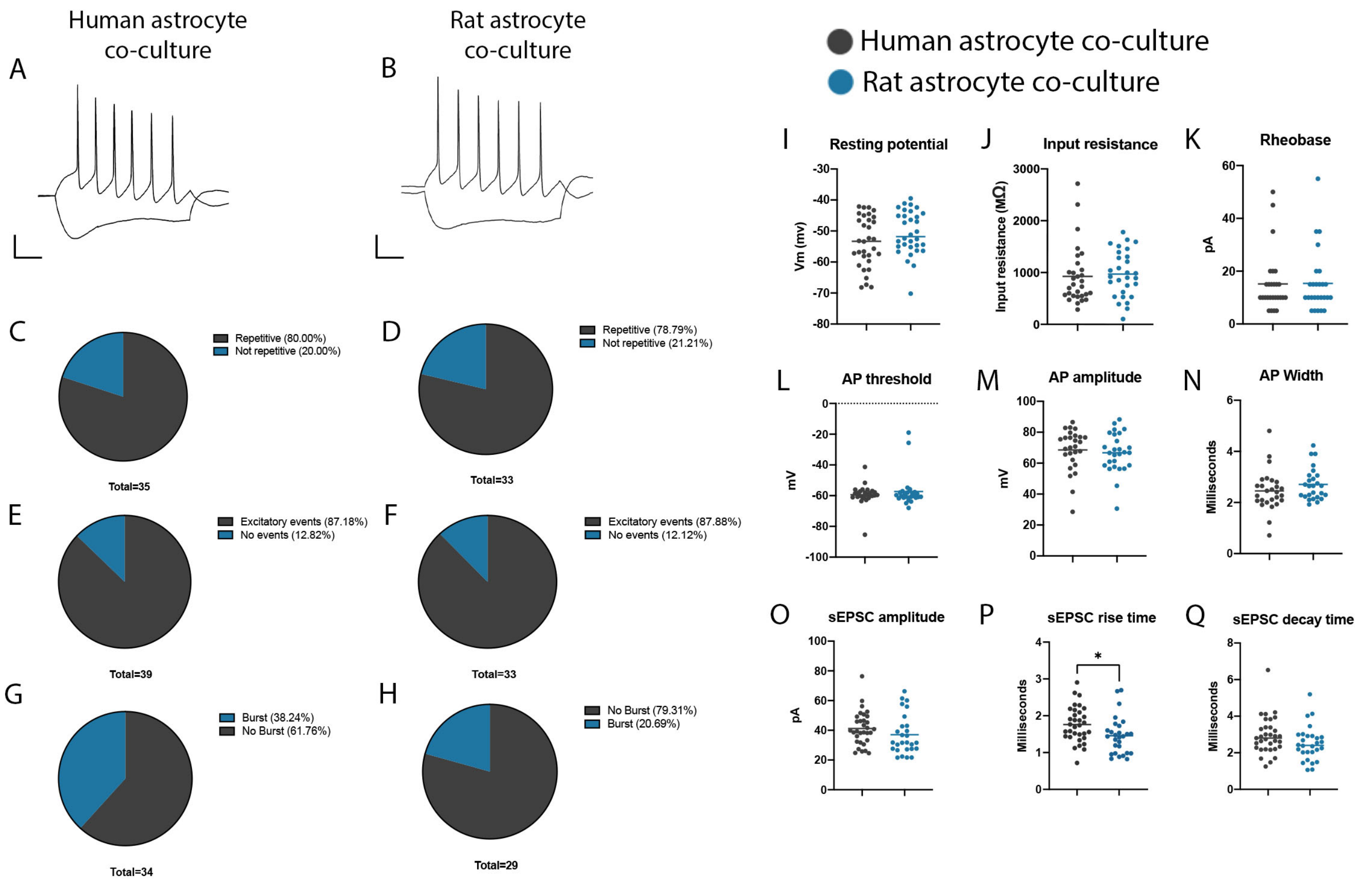


**Supplementary Figure 6: Whole-cell electrophysiological recordings of two-week old neuronal co-cultures with human or rodent astrocytes.** (**A**), (**B**) Representative traces of evoked action potentials in cultures with human (A) or rat (B) astrocytes. (**C**), (**D**) Percentage of neurons able to fire repetitive action potentials upon current injection in cultures with human (C) or rat (D) astrocytes. (**E**), (**F**) Percentage of neurons that receive spontaneous synaptic input in cultures with human (E) or rat (F) astrocytes. (**G**), (**H**) Percentage of neurons that received bursts of post synaptic currents in cultures with human (G) or rat (H) astrocytes, this percentage was non-significantly increased in cultures with human astrocytes. (**I**) – (**N**) Resting membrane potential (I), input resistance (J), rheobase (K), AP threshold (L), AP amplitude (M) and AP width (N) were similar in co-cultures with human (black) or rat (blue) astrocytes (n= 29 (human) and 30 (rat) cells). (**O**) sEPSC amplitude was similar in both conditions. (**P**) sEPSC rise time was slower in co-cultures with human astrocytes (two-tailed t-test, $P<0.05$). (**Q**) No differences were found in the decay time of sEPSC (n= 27 (human) and 32 (rat) cells).

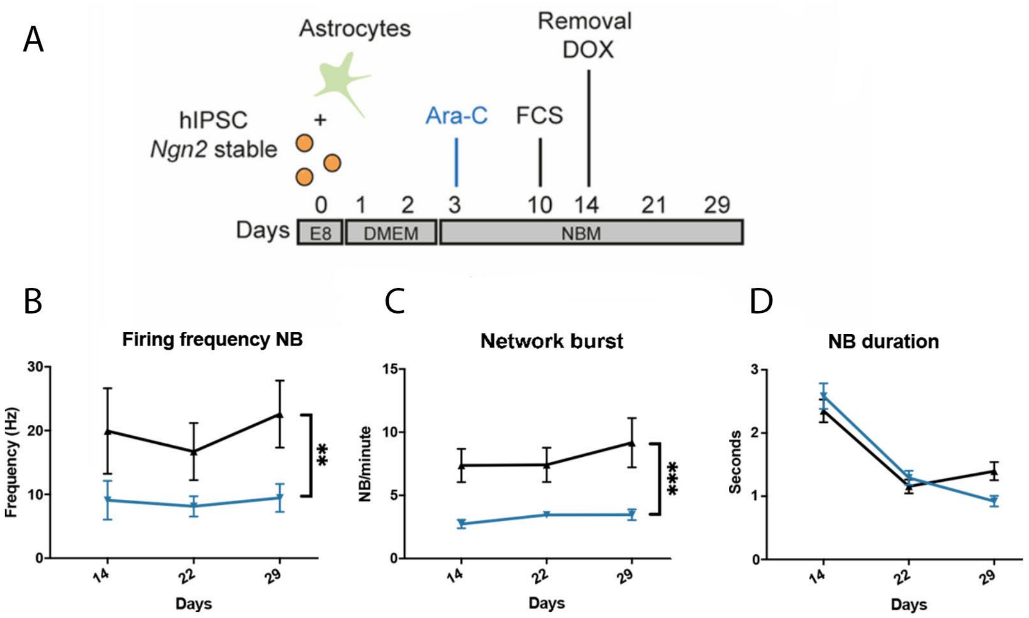


**Supplementary Figure 7: Implementation of hPSC-derived astrocytes in an independent laboratory.** (**A**) Experimental setup of neuronal co-culture. iPSCs are plated together with astrocytes in a co-culture and *Ngn2* overexpression is induced in iPSCs using doxycycline to initiate neuronal differentiation. (**B, C, D**) Analysis of *Ngn2*-neuronal co-cultures. Mean firing frequency (B) within NBs and NB rate per minute (C) are increased in human astrocyte co-cultures, while network burst duration is similar across conditions (D).

| Sample Clusters | Co-culture astrocytes | Mono-culture astrocytes |
|---|---|---|
| 1 - Astrocyte 2 | 1075 | 484 |
| 2 - Dividing | 289 | 1030 |
| 3 - Radial glia 2 | 521 | 548 |
| 4 - Dividing | 325 | 704 |
| 5 - Radial glia 2 | 150 | 701 |
| 6 - Radial glia 2 | 126 | 684 |
| 7 - Dividing | 85 | 456 |
| 8 - Radial glia 2 | 288 | 213 |
| 9 - Radial glia 2 | 31 | 338 |
| Total cells | 2890 | 5158 |

**Supplementary Table 1:** Cluster composition of integrated astrocyte sample

| Sample Clusters | Co-culture neurons | Mono-culture Neurons |
|---|---|---|
| 1 - Excitatory neuron 1 | 1086 | 682 |
| 2 - Excitatory neuron 4 | 1025 | 570 |
| 3 - Excitatory neuron 4 | 663 | 585 |
| 4 - Excitatory neuron 4 | 715 | 351 |
| 5 - Excitatory neuron 4 | 429 | 602 |
| 6 - Excitatory neuron 4 | 326 | 415 |
| 7 - Excitatory neuron 4 | 505 | 136 |
| 8 - Excitatory neuron 4 | 237 | 206 |
| 9 - Excitatory neuron 4 | 173 | 108 |
| 10 - Excitatory neuron 4 | 51 | 55 |
| 11 - Excitatory neuron 1 | 76 | 14 |
| 12 - Excitatory neuron 4 | 67 | 22 |
| Total cells | 5353 | 3746 |

**Supplementary Table 2:** Cluster composition of integrated neuron sample

**Supplementary Table 3:** Full list of differentially expressed genes in mono- and co-culture astrocytes

**Supplementary Table 4:** Full list of differentially expressed genes in mono- and co-culture *Ngn2*-neurons

| Gene | Expression in mono-culture astrocytes | Expression in co-culture astrocytes | Log2fold change | Adjusted p-value |
|---|---|---|---|---|
| **Top astrocyte progenitor genes** | | | | |
| *HIST1H2AI* | 0.00 | 0.01 | 0.01 | 0.21 |
| *HIST1H3E* | 0.03 | 0.02 | -0.02 | 1.00 |
| *HIST1H3B* | 0.34 | 0.11 | -0.23 | >0.01 |
| *HIST1H1B* | 0.66 | 0.23 | -0.43 | >0.01 |
| *PPDPF* | 1.54 | 2.97 | 1.44 | >0.01 |
| *TPX2* | 1.58 | 1.00 | -0.58 | >0.01 |
| *NUSAP1* | 1.45 | 1.08 | -0.37 | >0.01 |
| *HIST1H2AC* | 0.27 | 0.26 | -0.03 | 1.00 |
| *HIST2H2AC* | 0.57 | 0.21 | -0.36 | >0.01 |
| *TNC* | 0.34 | 0.38 | -0.18 | 1.00 |
| *KIF15* | 0.95 | 1.15 | -0.19 | >0.01 |
| *HIST1H1A* | 0.01 | 0.00 | 0.00 | 1.00 |
| *HIST1H2BC* | 0.02 | 0.12 | -0.14 | >0.01 |
| *FABP5* | 0.03 | 0.076 | 1.21 | >0.01 |
| *DTYMK* | 1.14 | 0.83 | -0.32 | >0.01 |
| *RAB11B* | 0.08 | 0.24 | 0.16 | >0.01 |
| *HES6* | 1.40 | 0.67 | 0.73 | >0.01 |
| *LRIG3* | 0.13 | 0.10 | 0.05 | 1.00 |
| *E2F5* | 0.30 | 0.31 | 0.04 | 1.00 |
| *MPPED2* | 0.02 | 0.05 | 0.03 | >0.01 |

| Gene | Expression in mono-culture astrocytes | Expression in co-culture astrocytes | Log2fold change | Adjusted p-value |
|---|---|---|---|---|
| **Top mature astrocyte genes** | | | | |
| *AGXT2L1* | 0.00 | 0.00 | 0.00 | 1.00 |
| *S100A1* | 0.00 | 0.01 | 0.01 | 1.00 |
| *SLC14A1* | 0.00 | 0.00 | 0.00 | 1.00 |
| *TMEM176A* | 0.00 | 0.00 | 0.00 | 1.00 |
| *TMX2* | 0.44 | 0.72 | 0.28 | >0.01 |
| *HHATL* | 0.00 | 0.00 | 0.00 | 1.00 |
| *PADI2* | 0.01 | 0.00 | 0.01 | 1.00 |
| *TLR4* | 0.02 | 0.01 | 0.02 | 0.96 |
| *HSD17B6* | 0.01 | 0.05 | 0.04 | >0.01 |
| *CHI3L1* | 0.01 | 0.01 | 0.04 | 1.00 |
| *NUDT3* | 0.95 | 1.15 | 0.19 | >0.01 |
| *FBXO2* | 0.02 | 0.12 | 0.14 | >0.01 |
| *NTSR2* | Not detected | Not detected | - | - |
| *ALDH1L1* | 0.03 | 0.076 | 0.07 | >0.01 |
| *ALDOC* | 0.01 | 0.039 | 0.04 | >0.01 |
| *SLC1A2* | 0.00 | 0.05 | 0.06 | >0.01 |
| *RYR3* | 0.10 | 0.02 | -0.08 | >0.01 |
| *GABRA2* | 0.00 | 0.03 | 0.03 | >0.01 |
| *CPE* | 0.75 | 1.26 | 0.51 | >0.01 |
| *GLUL* | 0.33 | 0.58 | 0.50 | >0.01 |

**Supplementary Table 5:** Top differentially expressed genes in human astrocyte progenitor cells and mature astrocytes as previously reported by Zhang et al[8]. Highlighted genes are significantly (Bonferroni corrected) up- (green) or downregulated (red) in iPSC-derived astrocytes when grown in a co-culture with Ngn2-neurons.
